# The transcription factor ZNF469 regulates collagen production in liver fibrosis

**DOI:** 10.1101/2024.04.25.591188

**Authors:** Sebastian Steinhauser, David Estoppey, Dennis P. Buehler, Yanhua Xiong, Nicolas Pizzato, Amandine Rietsch, Fabian Wu, Nelly Leroy, Tiffany Wunderlin, Isabelle Claerr, Philipp Tropberger, Miriam Müller, Lindsay M. Davison, Quanhu Sheng, Sebastian Bergling, Sophia Wild, Pierre Moulin, Jiancong Liang, Wayne J. English, Brandon Williams, Judith Knehr, Marc Altorfer, Alejandro Reyes, Craig Mickanin, Dominic Hoepfner, Florian Nigsch, Mathias Frederiksen, Charles R. Flynn, Barna D. Fodor, Jonathan D. Brown, Christian Kolter

**Author notes:** Co-senior authors (Charles R. Flynn, Barna D. Fodor, Jonathan D. Brown, Christian Kolter). All authors except for LMD, QS, DPB, JL, WJE, BW, JDB, CRF, are or were employees of Novartis Pharma. LMD, QS, DPB, JL, WJE, BW, JDB, and CRF declare no conflict of interest exists. PM is chief scientific officer at Deciphex Ltd, Dublin, Ireland.

## Abstract

Non-alcoholic fatty liver disease (NAFLD) - characterized by excess accumulation of fat in the liver - now affects one third of the world’s population. As NAFLD progresses, extracellular matrix components including collagen accumulate in the liver causing tissue fibrosis, a major determinant of disease severity and mortality. To identify transcriptional regulators of fibrosis, we computationally inferred the activity of transcription factors (TFs) relevant to fibrosis by profiling the matched transcriptomes and epigenomes of 108 human liver biopsies from a deeply-characterized cohort of patients spanning the full histopathologic spectrum of NAFLD. CRISPR-based genetic knockout of the top 100 TFs identified ZNF469 as a regulator of collagen expression in primary human hepatic stellate cells (HSCs). Gain- and loss-of-function studies established that ZNF469 regulates collagen genes and genes involved in matrix homeostasis through direct binding to gene bodies and regulatory elements. By integrating multiomic large-scale profiling of human biopsies with extensive experimental validation we demonstrate that ZNF469 is a transcriptional regulator of collagen in HSCs. Overall, these data nominate ZNF469 as a previously unrecognized determinant of NAFLD-associated liver fibrosis.

## INTRODUCTION

Non-alcoholic fatty liver disease (NAFLD) is the most common form of liver disease in the developed world (*1*). The condition is closely associated with obesity and insulin resistance often due to lifestyle factors such as high-calorie diets and sedentary behavior. NAFLD exists across a spectrum of clinical and pathologic stages characterized by 1) steatosis, 2) inflammation (a.k.a steatohepatitis) and 3) fibrosis/cirrhosis (*2*). Morbidity and mortality from NAFLD significantly escalate during the transition to fibrosis/cirrhosis. Despite the concerning epidemiologic trends for NAFLD (*3*), only one drug therapy has been recently approved for NASH with moderate to advanced fibrosis (*4*). Orthotopic liver transplantation is the only curative treatment for late stage NAFLD patients. Notably, reversal of liver fibrosis has been achieved by bariatric surgery (*5*) and in patients with chronic Hepatitis B and C in response to long-term eradication of viruses with targeted anti-viral therapies (*6*)(*7, 8*). As such, the potential to forestall or reverse this morbid and fatal stage of disease renders liver fibrosis an intently pursued therapeutic target in NAFLD.

Activated hepatic stellate cells (aHSCs) are a key cell-type that drives fibrogenesis in all forms of chronic liver disease, including NAFLD (*9*). HSCs comprise approximately 5-10% of the non-parenchymal cell population in the liver (*10*). In physiologic contexts, resident HSCs exist in a quiescent state (qHSCs), serving as a storage depot for vitamin A (*11*). However, in response to toxins, inflammatory signals and growth factors, HSCs become activated and assume a myofibroblast-like cell state capable of migration, proliferation and collagen production (*12*).

Reversible changes in cell state are orchestrated by an interplay between multiple transcription factors (TFs) and chromatin-dependent signaling pathways. In HSCs, the overall balance of activity between lineage-determining TFs (LDTFs) along with signal-responsive TFs (SRTFs) determines the inflammatory and fibrogenic gene expression programs (*13–17*).

More recently, occupancy maps of chromatin coactivators coupled with genome-wide enhancer profiles - e.g. measured by acetylation of histone 3 at lysine 27 (H3K27ac, a histone mark of active regulatory regions) - have been used to reconstruct transcriptional circuits involving a core set of TFs that orchestrate cell-type specific gene expression programs (*18–21*). Whether this conceptual framework could be used to discover TFs that drive disease-specific cell state transitions in human NAFLD has never been tested. The identification and therapeutic targeting of transcriptional regulators involved in stage-specific disease progression could reveal new approaches to treat fibrotic NASH.

In this study, we integrate RNA sequencing (RNA-seq) with enhancer profiles of liver tissue obtained from obese patients at the time of bariatric surgery to identify candidate TFs predicted to regulate collagen expression and fibrosis. A CRISPR knockout screen of these prioritized TFs in human HSCs reveals *ZNF469* - a causal gene in Brittle Cornea Syndrome featuring abnormal extracellular matrix production - as a regulator of subsets of collagen genes and genes involved in extracellular matrix (ECM) homeostasis. The specificity in regulation of collagen reflects direct ZNF469 binding to the genes of collagen types 1, 3 and 5. Regulation of ZNF469 is conserved in mouse stellate cells and murine models of liver fibrosis and NAFLD as well as other human fibrotic diseases. Collectively, our data reveal how integration of multiomics readouts can be leveraged to gain an improved understanding of transcriptional regulators involved in the pathogenesis of HSC activation and liver fibrosis.

## RESULTS

### Integration of transcriptomics and cis-regulatory landscapes in human NAFLD livers predicts activity of TFs involved in fibrosis

To extract gene signatures in human NAFLD we characterized the transcriptomes of wedge liver biopsies collected from subjects at the time of bariatric surgery. Thirty patients with NAFL, 50 patients with NASH and 28 patients without liver disease (histologically normal) were identified using NASH CRN scoring criteria by a pathologist blinded to the clinical data (fig. S1C). Among these 108 patients the mean age was 43 years, BMI was 46.6 kg/m^2^, 84.3% were women, 28.7% had type 2 diabetes, 21.2% had hypercholesterolemia, 48.1% had high blood pressure and 26.8% were current or former smokers. Regarding medication use, 33% were on a PPI, 28% on a SSRI, 22% on HRT and 26% on a NSAID, table S1) Using this deeply characterized cohort, we identified 582 upregulated and 143 downregulated genes when comparing NASH patients with a fibrosis score of 2 or 3 (NASH_F2/F3) to healthy (NOR) liver tissue (FDR <= 0.01 & |log2FC| > 0.5, Fig. 1A, table S2).

**Figure 1.**
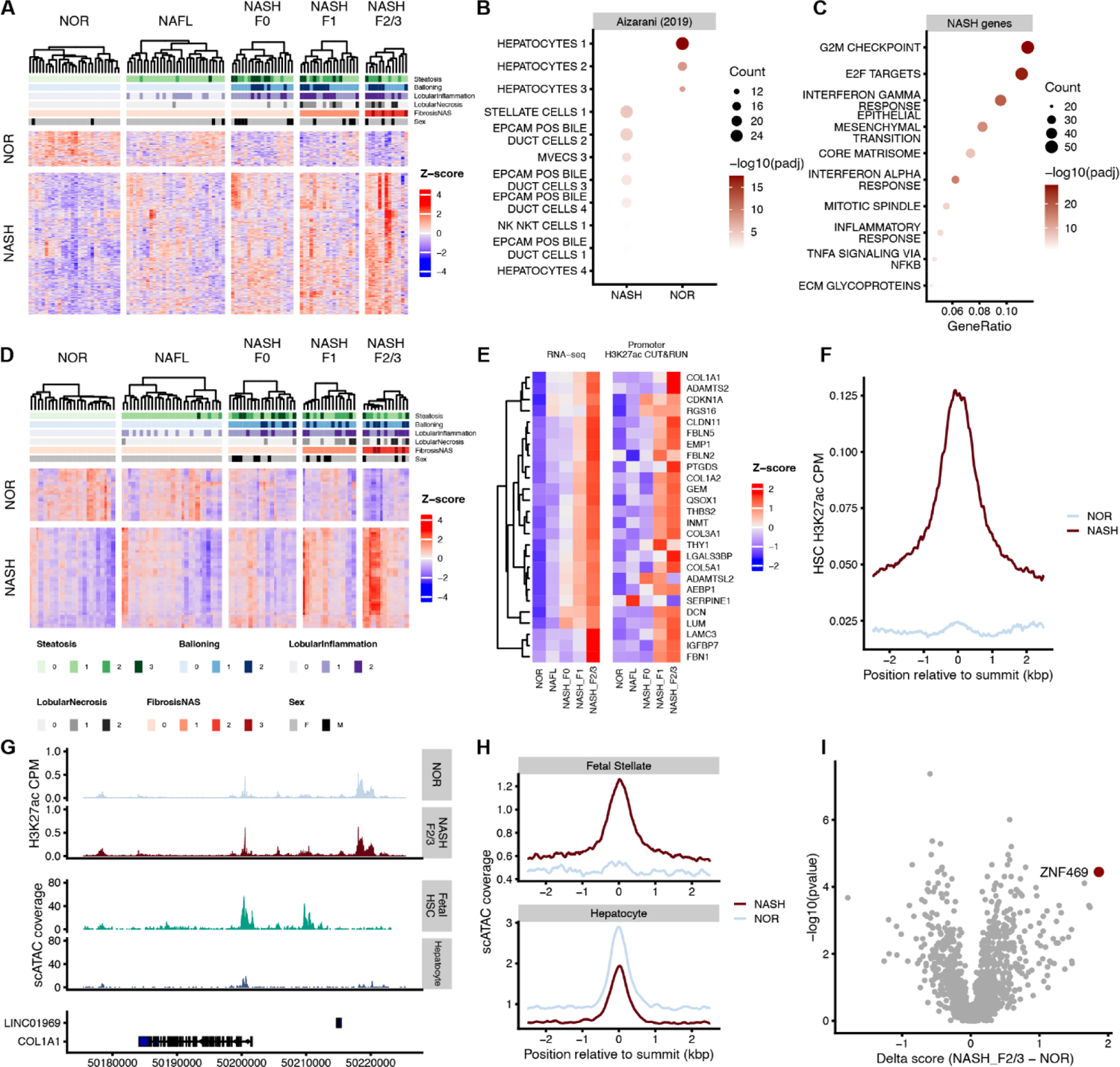
Integration of transcriptomics and cis-regulatory landscapes in human NAFLD livers predicts activity of TFs involved in fibrosis. (**A**) Heatmap of RNAseq data normal vs NASH. (**B**) Enrichment plot of cell types based on RNAseq, (**C**) Enrichment plot of GO terms, (**D**) Heatmap of CUT&RUN H3K27ac data clustered by normal vs NASH. (**E**) Heatmap of expression across the top enriched genes in NASH vs. normal concatenated by H3K27ac change at promoters. (**F**) Correlation of H3K27ac from primary HSC compared to H3K27ac from the cohort. (**G**) Genome browser view of the H3K27ac profile at the *COL1A1* locus compared to publicly available scATAC-seq data from cultured fetal HSCs and hepatocytes. (**H**) Correlation plot comparing scATAC-seq signal in HSCs or hepatocytes and H3K27ac signal in the NAFLD cohort. (**I**) Volcano plot of predicted TF activity generated with DoRothEA analysis comparing NASH_F2/3 vs. Normal (NOR) samples.

Having identified these signature genes between NOR and NASH_F2/F3 we next performed a gene set enrichment analysis on publicly available scRNA-seq data to ascertain whether these signatures were enriched in specific cell types in the liver (*22*) (*14*). We observed an enrichment of hepatocyte marker in genes upregulated in healthy livers whereas HSC markers were enriched in genes upregulated in NASH livers (FDR < 0.01, Fig. 1B). Activated HSCs play a known and key role in hepatic fibrosis (*15*). Thus, our finding that the disease-enriched signature genes mapped back to HSCs is consistent with current paradigms of liver fibrosis (*17*). Based on this result, we then performed pathway analysis to determine whether these genes are functionally related. The genes upregulated in NASH are enriched in pathways related to epithelial-mesenchymal transition (EMT) and core matrisome biology, both consistent with an increase in fibrogenesis (Fig. 1C) and consistent with previously published work (*13, 16*).

### Inference of transcription factor activity from cis-regulatory landscapes in NAFLD

NAFLD is a complex disease with multiple signals integrated at the level of gene regulation and governed by the interplay between master regulatory TFs and chromatin signaling. Thus, we aimed to infer activity of TFs between NASH_F2/3 disease stage and histologically normal livers (*36, 37*). We profiled genome-wide H3K27ac signal using “Cleavage Under Targets & Release Using Nuclease” (also known as CUT&RUN) in 99 liver samples from the VUMC biorepository (*36, 37*). Overall, our analysis identified 14,348 genome regions with differential H3K27ac abundance, when comparing the cases of NASH_F2/3 versus histologically normal livers (Fig. 1D; FDR <= 0.01 & |log2FC| > 0.5, table S3). We found that 4,937 regions were enriched in normal livers, while 9,411 regions were associated with disease (Fig. 1D). Moreover, 15 out of 26 genes that were most significantly regulated on a transcriptomic and chromatin level during the transition to fibrosis (NASH_F23 vs normal) were genes linked to collagen-containing extracellular matrix (e.g. *COL1A1, COL1A2, COL3A1, COL5A1, IGFBP7, DCN, AEBP1, LGALS3BP, THBS2, FBLN2, FBLN5, FBN1, SERPINE1, ADAMTS2*, and *LUM*; Fig. 1E). Notably, genomic regions with enriched H3K27ac signals in NASH displayed high levels of H3K27ac signal also in HSC monocultures *in vitro* (Fig. 1F and fig. S1D), suggesting a contribution of HSCs in the epigenetic landscape rewiring in disease development and progression. To confirm this observation, we compared our NOR and NASH_F2/3 regions to scATAC-seq data from Zhang *et al* to identify cell type specific regulatory regions. We identified disease-specific peaks of H3K27ac in the promoter as well as upstream intergenic regions of the gene *COL1A1*, a core extracellular matrix component associated with fibrosis, that overlapped with scATAC peaks from HSCs (Fig. 1G and fig. S1E).

We could further show that HSC-enriched ATAC-seq signals displayed higher levels of chromatin accessibility in NASH biopsies compared to normal biopsies (Fig. 1H). These results suggest that gene expression together with H3K27ac profiling capture distinct disease and normal cell states within the entire cohort.

The data demonstrate that patterns of enhancer signal and gene expression can identify disease stages, which suggests that we could use this information to infer transcription factor activity. To test this concept, we used 6 different computational tools: ANANSE (*23*), CRCmapper (*24*), DoRothEA (*25*), RcisTarget (*26*), MonaLisa (*27*), and HOMER (*28*) to call TFs that might explain the chromatin and transcription changes in diseased livers by pairwise comparison between normal and NASH_F2/F3 samples. In total, those tools identified 684 TFs and more than 50% (369 TFs) were identified only with one method. For instance, ZNF469 showed the highest activity at NASH_F2/3 compared to normal samples according to DoRothEA (Fig. 1I) but was not identified by any of the other methods (table S4).

### CRISPR loss-of-function screen identifies transcriptional regulators of collagen production in human HSCs

To validate our findings experimentally we performed an arrayed, CRISPR loss-of-function screen focusing on 100 TFs that were selected by multiple criteria (e.g. absolute score per method, hit in multiple tools, expression in HSCs, table S5-7). Given the strong links to HSCs as determined by the transcriptomics as and enhancer profiling in patient tissues, we performed the CRISPR assays in primary human HSCs (Fig. 2A). We quantified, at single-cell resolution, the difference of intracellular COL1A1 between the crRNA-tracrRNA complexes (for simplicity these complexes will be referred to as “sgRNA”) targeting each TF compared to the non-targeting sgRNA (hNTO) controls. We identified 18 TFs that downregulated COL1A1 (Fig. 2B-D) compared to the median COL1A1 expression of the hNTO (threshold < −200 in Fig. 2B). Since cell death could also decrease intracellular COL1A1, the number of cells was used to determine viability. We observed that our cell viability controls (such as POLR2B) and predicted or previously identified essential TFs in myofibroblasts, such as MYC and FOXM1 (*29*), decreased cell viability and were discarded from further experiments (fig. S2A). Therefore, we nominated 11 candidate TFs for further validation, including *ZNF469* as the top hit.

**Figure 2.**
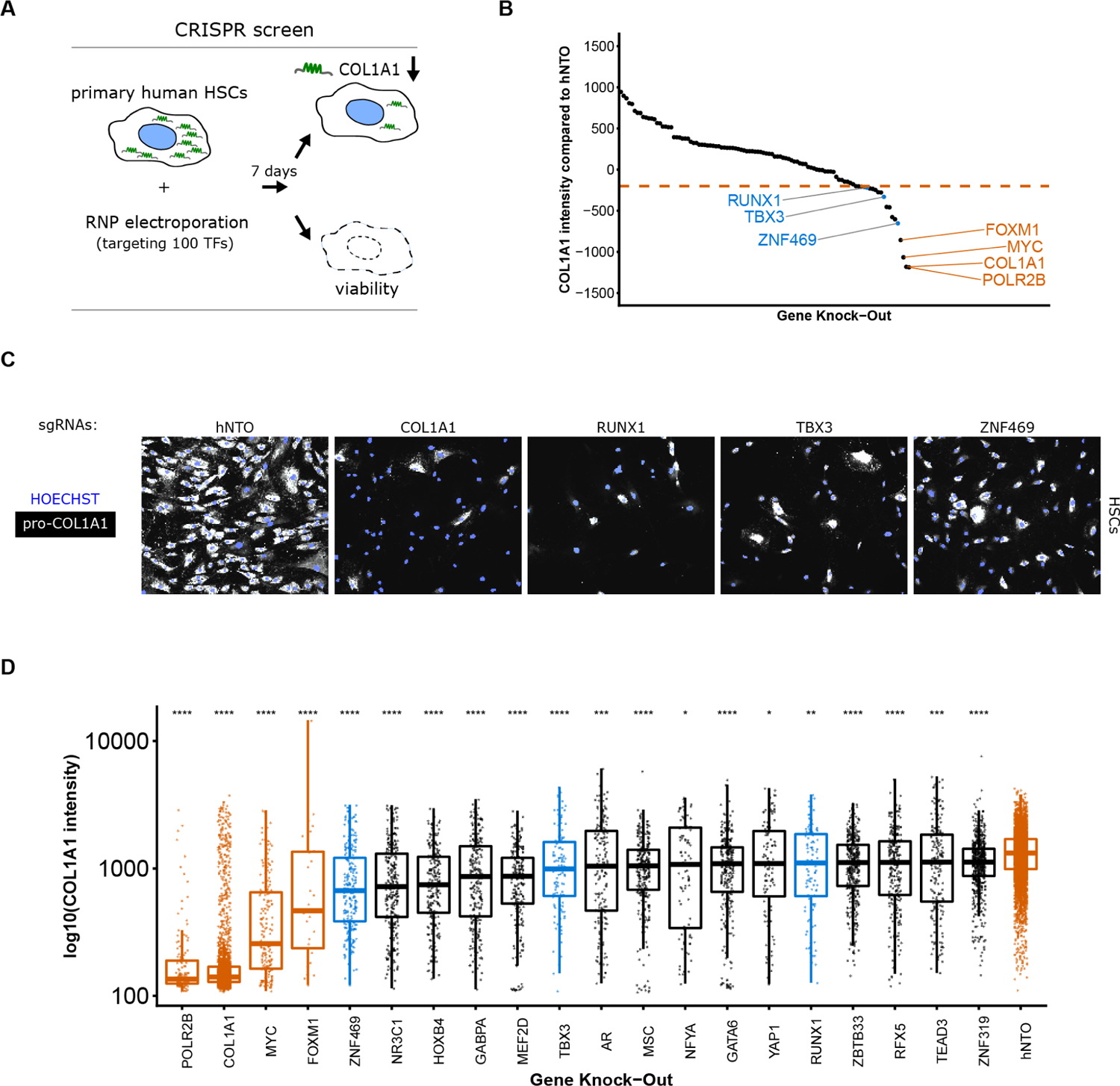
CRISPR loss-of-function screen identifies transcriptional regulators of collagen production in primary human HSCs. **(A)** Schematic of targeted CRISPR screen with single-cell high-content imaging read-out of collagen protein expression. **(B)** Waterfall plot of primary screen results at day7. **(C)**Representative IF photomicrographs showing COL1A1 staining in the following CRISPR KO conditions: non target control (hNTO), *COL1A1* KO, *RUNX1* KO, *TBX3* KO and *ZNF469* KO. **(D)** Box plots showing distribution of COL1A1 expression by IF upon KO of top TF hits, each dot corresponds to COL1A1 IF measurement in one cell. We used a Wilcoxon test to compare the medians of the COL1A1 immunofluorescence expression. (* = P ≤ 0.05; ** = P ≤ 0.01, *** = P ≤ 0.001, **** = P ≤ 0.0001).

### ZNF469 knockout alters collagen mRNA expression in HSCs

To further prioritize the top candidates, we performed RNA-seq experiments in order to: 1) filter out factors that regulate COL1A1 protein levels post-transcriptionally, 2) measure the specificity of the transcriptional effects on the whole gene signature compared to global effects, and 3) control CRISPR editing efficiency of each target (fig. S3A and S3B). The latter point is particularly important for genes such as *ZNF469,* as its transcript harbors one large coding exon and CRISPR mutations do not lead to nonsense mediated mRNA decay as previously demonstrated for the mouse homolog (*30*). Notably, only the targeting of three TFs (*ZNF469*, *RUNX1*, *TBX3*) resulted in a significant (FDR < 0.01) downregulation of *COL1A1* mRNA (Fig. 3A). All of these TF perturbations also downregulated *COL1A2,* located on a different chromosome from *COL1A1* (Fig. 3A) - suggesting that they are upstream regulators of type 1 collagens. Compared to *RUNX1* KO and *ZNF469* KO, *TBX3* KO had the smallest effect on *COL1A1* and *COL1A2* mRNA (Fig. 3B). When considered globally, similar gene sets were enriched for *ZNF469*, *RUNX1* and *TBX3* KO including extracellular matrix (ECM) organization (Fig. 3C and fig. S3C). In particular, for *RUNX1* KO the two top significantly downregulated genes were *LUM* and *THBS2*. For *ZNF469* KO the two most significantly downregulated genes were *COL1A1* and *COL1A2* and we confirmed this in a second donor (Fig. 3B and fig. S3E). All four genes (*LUM*, *THBS2*, *COL1A1* and *COL1A2*) are part of the collagen-containing ECM GO term (GO: 0062023). As the disease-specific upregulated transcripts in liver samples also contained many ECM genes, we explored whether these TFs reduce expression of genes in the signature, restricting our analysis to targets that were expressed in HSCs (Fig 1F). While both *TBX3* and *RUNX1* showed a strong downregulation of most of these genes, *ZNF469* KO had a more specific effect on *COL1A1*, *COL1A2* and *COL3A1* as well as *INMT*, *ADAMTSL2* and *PTGDS* (Fig. 3D and fig. S3D). In addition to this HSC-specific panel, *ZNF469* KO also significantly downregulated *FMOD* and *FNDC1*, two other fibrosis-related genes (Fig. 3B) (*31*).

**Figure 3.**
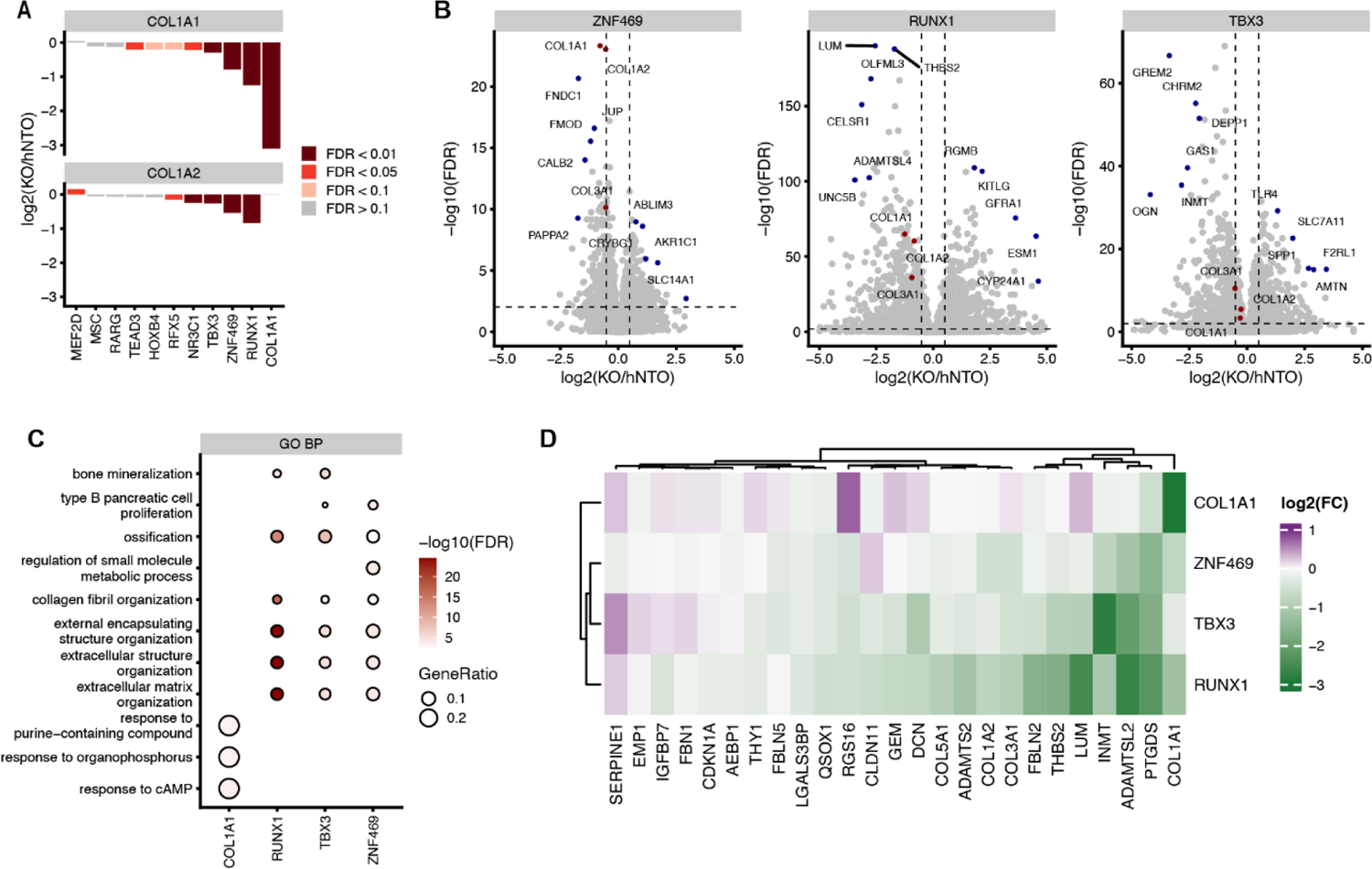
ZNF469 knockout alters collagen mRNA expression in HSCs. **(A)** Bar plot of *COL1A1* and *COL1A2* gene expression obtained by RNAseq, after KO of top hits from the CRISPR Screen: *MEF2D, MSC; RARG, RFX5, HOXB4, TEAD3, NR3C1, TBX3, ZNF469, RUNX1*. **(B)** RNAseq volcano plots for *ZNF469*, *TBX3* and *RUNX1* with *COL1A1* and *COL1A2* highlighted. **(C)** Gene set enrichment of downregulated genes upon *ZNF469*, *RUNX1* or *TBX3* KO. **(D)** Heatmap of differential HSC genes (between NASH_F2/3 and normal) from Fig. 1F upon CRISPR KO with z-score distribution across hNTO, *COL1A1*, *ZNF469*, *TBX3* and *RUNX1*.

### ZNF469 localizes to gene bodies of collagen genes in HSCs

The CRISPR screen, as well as mRNA profiling suggest that ZNF469 plays a selective role in the regulation of collagen gene expression. However, there is a lack of functional data demonstrating that it is a *bona fide* DNA binding protein. Thus, we chose to explore its biological role in HSCs more fully. We annotated its sequence with previously reported- and protein-sequence-analysis-based features which were in agreement with putative TF-, nucleic acid- and protein-binding functions (Fig. 4A) (*32–35*). To address questions about ZNF469 function, we designed a doxycycline-inducible piggyBac transposon system to express the full-length gene with an N-terminal 3xHA epitope tag (HA-ZNF469) and another construct with a deletion harboring 6 of the 8 zinc fingers (referencing this region in the rest of the manuscript as “zinc finger domain” for simplicity) (del:aa3071-3727; HA-Δzf-ZNF469) (Fig. 4A). For practical reasons we turned to the LX-2 cell line, which is an immortalized human stellate cell line that recapitulates features of the activated HSC. Immunohistochemistry results with an anti-ZNF469 antibody indicated that ZNF469 localized exclusively to the nucleus in LX-2 cells after doxycycline treatment (Fig. 4B). Motivated by this result, we next mapped the genome-wide binding sites of endogenous ZNF469 in HSCs using CUT&RUN with the same antibody and in the transgenic LX-2 cells. Cistromic analyses demonstrated a specific gene-body distribution of ZNF469 at collagen genes, including *COL1A1* (Fig. 4C), *COL1A2* and *COL3A1* (fig. S4A). In particular, ZNF469 was bound to the promoter of *COL1A1* and several putative enhancer elements in proximity to this gene. No DNA binding of ZNF469 was observed at these loci upon KO of endogenous ZNF469 in HSC. Moreover, LX-2 cells expressing the deletion construct also did not show any DNA binding providing important controls for the antibody and CUT&RUN overall (Fig.4C and fig. S4A). By differential analysis of chromatin profiles in control cells vs. ZNF469 KO cells, ZNF469 signal was significantly higher at intragenic sites including transcriptional start sites of collagen genes (Fig. 4D and E).

**Figure 4.**
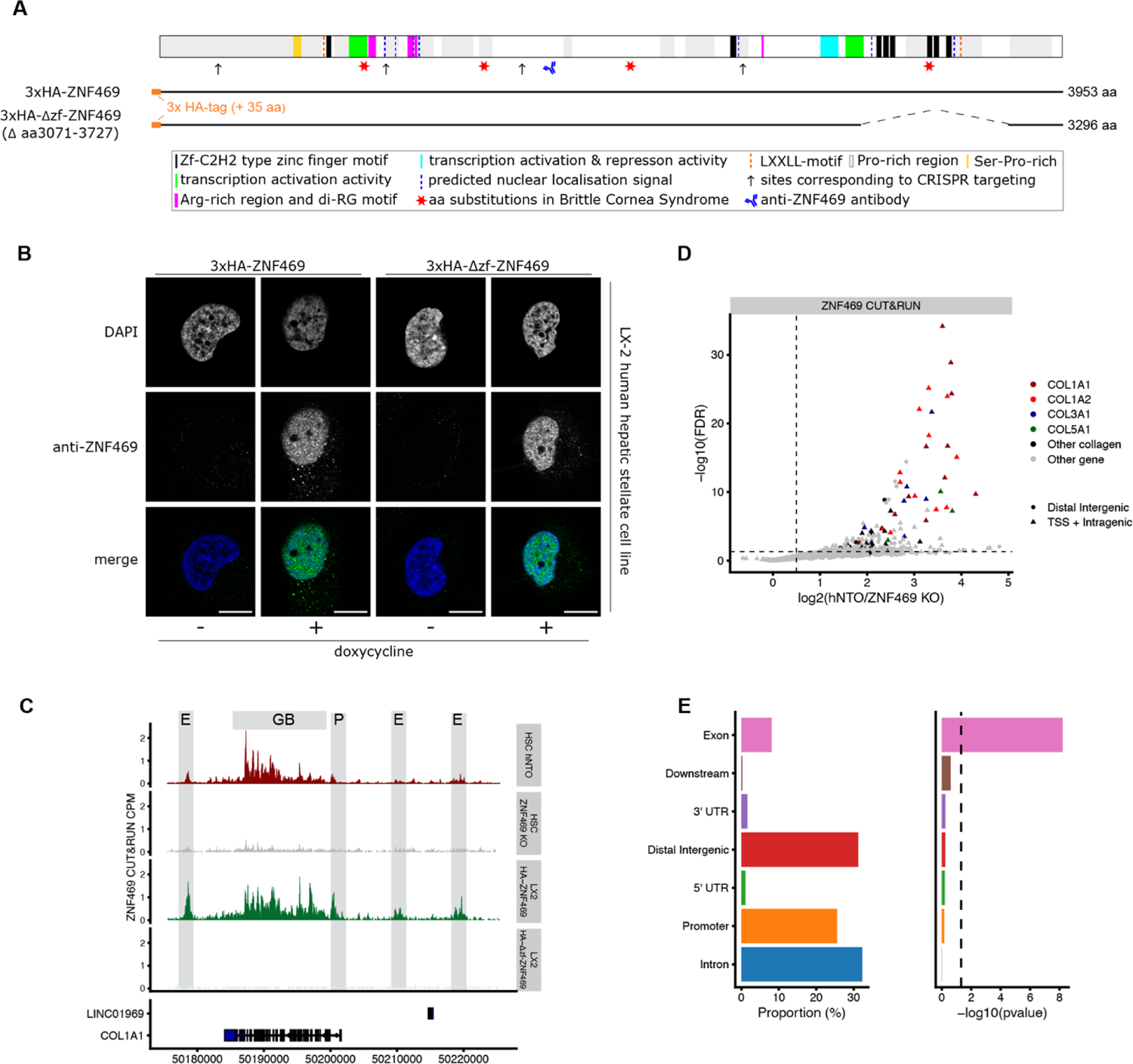
ZNF469 is enriched at extracellular matrix related genomic sites. **(A)** Schematic of ZNF469 protein (NP_001354553.1) with previously reported and predicted sequence features, regions active in transactivation domain reporter assays, aa substitutions linked to the Brittle Cornea Syndrome, anti-ZNF469 antibody antigen region, and positions corresponding to the CRISPR sgRNA target sites. Coverage of ZNF469 by full length and deletion harboring cDNA constructs used for stable inducible cell line generation are shown below (see also table S8). **(B)** representative photomicrographs of ZNF469 subcellular localization by indirect immunofluorescence staining with the anti-ZNF469 antibody in transgenic LX-2 overexpressing cells; scale bar = 10 mM). **(C)** Genome browser tracks at the *COL1A1* locus of ZNF469 CUT&RUN signals generated with the anti-ZNF469 antibody in CRISPR experiments in none-targeting and *ZNF469* targeted human **HSCs** as well as transgenic LX-2 cells with doxycycline inducible full length or deletion harboring *ZNF469* cDNA (E: putative enhancer, P: promoter, GB: gene body) **(D)** Volcano plot of differential occupancy of endogenous ZNF469 in control cells vs. ZNF469 KO cells. Data shows change in signal at distal elements (circles) and TSS / intragenic sites (triangles) with color coded for collagen genes. (**E**) Bar plots of distribution of ZNF469 peaks at genomic annotations.

We next tested whether ZNF469 overexpression could rescue the *COL1A1* reduction when knocking out the endogenous *ZNF469* gene. For these series of experiments, we first generated new mouse stellate cell lines (JS1) targeting the mouse ortholog *Zfp469* using CRISPR guides against either the proximal portion of the gene or a region encoding for the ZF domain near the 3’-end of the gene. Both CRISPR treatments resulted in a ∼50% reduction in *Col1a1* and *Col1a2* mRNA in these cells as compared to non-targeting control (fig. S4B). Introduction of full-length HA-tagged human ZNF469 rescued the *Col1a1* and *Col1a2* mRNA levels and even increased *Col1a1* mRNA levels 2-4 fold compared to baseline levels. In addition, an orthogonal approach using transient siRNA knockdown of *Zfp469* resulted in ∼50% reduction in *Col1a1* and *Col1a2* mRNA when compared to a scrambled siRNA control (fig. S4C). These results suggest conservation of *ZNF469/Zfp469* as a regulator of collagen expression between human and mouse species.

### *ZNF469* knockout alters local chromatin structure at collagen loci in HSCs

To further confirm ZNF469 as a regulator of collagen expression, we profiled genome-wide H3K27ac following *ZNF469* CRISPR KO (Fig. 5A). *ZNF469* KO resulted in 171 differential H3K27ac peaks (FDR <= 0.01 & |log2FC| > 0.5). As shown with *ZNF469* binding, the greatest reductions in H3K27ac were observed at the *COL1A1* and *COL1A2* loci. In particular, the *COL1A2* locus was the most significantly differentially acetylated gene with multiple exons showing signals (Fig. 5A and fig. S5A). These results suggest that ZNF469 has a specific role in regulating collagen-type 1 genes. Notably, other collagens (e.g. *COL3A1, COL5A1)* were also amongst the top 10 affected loci (Fig. 5A). By integrating the RNA-seq and H3K27ac datasets, we observed that the downregulation of *COL1A1* and *COL1A2* mRNA correlates with a reduction of H3K27ac at those sites using promoter or enhancer annotation (Fig. 5C and fig. S5B). Of note, this pattern is the inverse of what was observed with the human liver samples in which ZNF469 was predicted to be more active and supports a directionally consistent link between ZNF469 action/expression and collagen expression (Fig. 1F). Since enhancer-dependent signaling can occur over long distances through chromatin looping to promoters, we next explored the 3-dimensional gene regulatory landscape in HSCs using MicroC-promoter capture. MicroC-promoter capture is a chromatin conformation assay that fragments DNA with micrococcal nuclease and then uses capture probes to pull down human promoters from the sequencing library. This approach allowed us to discover interactions between distal regulatory elements and human promoters in an unbiased way across large regions of the genome. Using wild type primary HSCs, we observed significant interactions of distal regions, likely enhancers, to the promoters of *COL1A1* and *COL1A2*, respectively (Fig. 5B-C, 5D upper panel and fig. S5A-B). Notably, a subset of these interactions overlapped with H3K27ac signals in human livers with NAFLD (Fig. 5D middle panel) and are altered in HSCs *in vitro* upon *ZNF469* KO (Fig. 5D lower panel).

**Figure 5.**
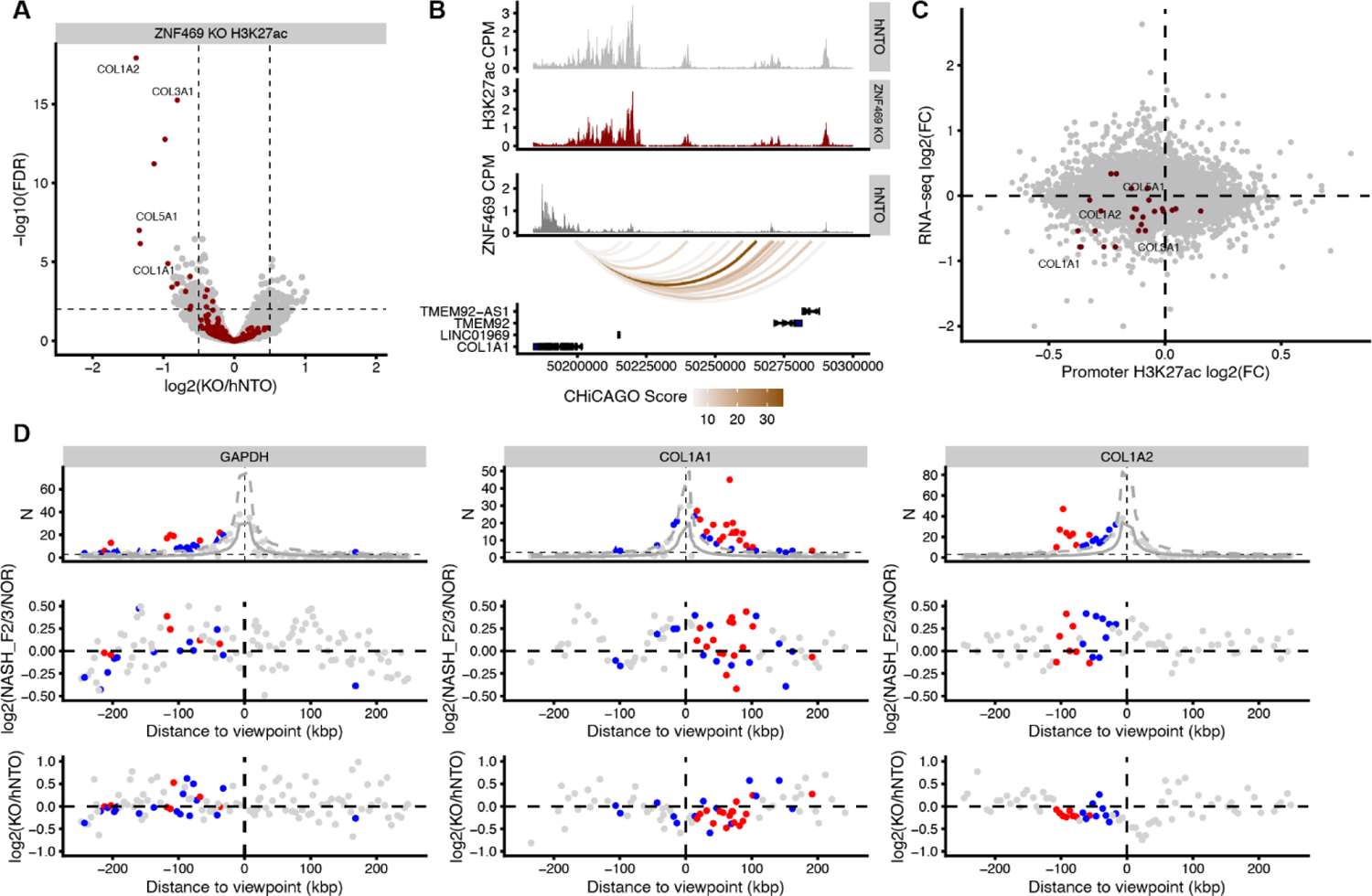
ZNF469 knockout alters local chromatin structure at collagen and ECM loci in HSCs. **(A)** Volcano plot of CUT&RUN H3K27ac with *COL1A1* and *COL1A2* highlighted upon *ZNF469 KO*. **(B)** Genome track of H3K27ac occupancy and promoter-capture-Micro-C at the *COL1A1* locus. **(C)** RNA-seq and CUT&RUN integration upon *ZNF469* KO highlighting *COL1A1* and *COL1A2* as the top affected genes. **(D)** 3D chromatin interactions. Top panel: promoter-capture-Micro-C data showing promoter bait plots for *GAPDH*, *COL1A1* and *COL1A2*. Interactions with a Chicago score ≥5 higher were considered significant (red dots); sub-threshold interactions (3 ≤ score < 5) are shown as blue dots. Grey lines show expected counts and dashed lines the upper bound of the 95 % confidence intervals; middle panel: H3K27ac signal of disease vs normal; colors refer to top panel; lower panel: H3K27ac human donor signal upon *ZNF469* KO, colors refer to top panel.

In summary, activity of enhancers at the *COL1A1* and *COL1A2* loci, as indicated by H3K27ac signal, were reduced following *ZNF469* KO, and chromatin conformation assays revealed likely interactions between these same regions and promoters of the collagen genes that are downregulated in response to *ZNF469* KO. Overall, these data provide further evidence that ZNF469 functions as a transcriptional regulator of collagen gene expression through enhancer-dependent signaling in HSCs.

### *ZNF469* expression correlates with fibrosis in human NAFLD

ZNF469 loss-of-function decreased both *COL1A1* and *COL1A2* expression, which encode two collagen proteins known to form heterotrimers. Thus, regulation of both genes by ZNF469 might modulate levels of fibrosis at the organ level. To explore this relationship, first we performed multiplexed RNA fluorescent *in situ* hybridization (RNA-FISH) in a subset of liver sections from the VUMC liver biorepository using MERSCOPE technology (Fig. 6A). For this assay, we designed a 300-gene probe set that included markers of aHSCs (e.g., *DCN*) and other cell types including hepatocytes (e.g. *HNF4a*) along with *COL1A1* and *ZNF469.* MERSCOPE enabled simultaneous measurements of the number and spatial distribution of *ZNF469* transcripts along with other cell marker genes in normal versus fibrotic NASH human livers (Fig. 6A-C, fig. S6A and table S9). We detected co-expression of *ZNF469* and *COL1A1, COL1A2* and *DCN* (Fig. 6D-E). Notably, we did not observe co-expression of ZNF469 with *HNF4A (*hepatocyte marker) or *MARCO* (Kupffer cell marker) (Fig. 6D). Hence, we demonstrated *ZNF469* transcript expression is localized in HSCs in human NAFLD livers with fibrosis from the VUMC repository.

**Figure 6.**
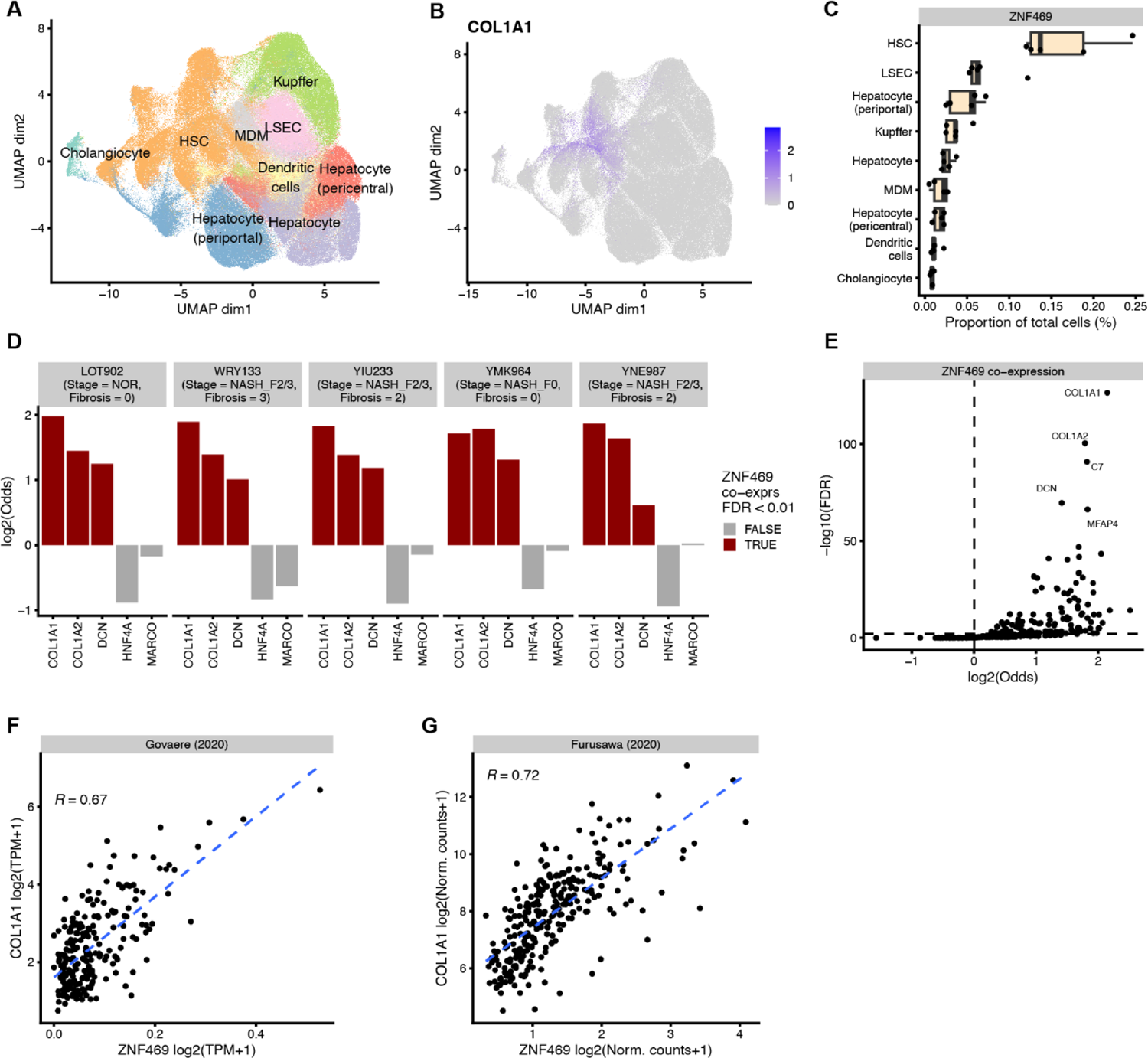
ZNF469 expression correlates with collagen production in HSCs in human NAFLD. (**A**) MERSCOPE UMAP showing liver cell types (**B**) colored by COL1A1 expression (**C**) *ZNF469* expression per identified cell type, (**D**) *ZNF469* is co-expressed with *COL1A1*, *COL1A2* and *DCN*, but not *HNF4α* and *MARCO,* (**E**) *COL1A1* and *COL1A2* are the top 2 co-expressed genes with *ZNF469*, (**F**) Scatter plot showing correlation (Pearson) of *ZNF469* and *COL1A1* expression in public NASH cohort. (**G**) Scatter plot showing correlation (Pearson) of *ZNF469* and *COL1A1* expression in published interstitial pulmonary fibrosis cohort.

We next wanted to explore whether the relationship between *ZNF469* expression and degree of fibrosis can be generalized using multiple additional clinical cohorts. Indeed, we observed in 5 out of 6 publicly available NAFLD as well as interstitial pulmonary fibrosis cohorts (IPF) a strong correlation (> 0.5) of *COL1A1* and *ZNF469* expression (Fig. 6F-G and fig. S6C-D).

Conservation of regulation is one way to determine potential biological relevance of predicted gene regulatory networks. We observed that *ZNF469/Zfp469* was significantly induced in two murine models of steatosis with fibrosis: Gubra Amylin (GAN) diet and choline-deficient, amino acid defined high fat diet (CDAHFD; fig. S7A). The increase in *Zfp469* was accompanied by an increase in *Col1a1* mRNA. We did not observe a difference in *Zfp469* and *Col1a1* expression in ethionine-treated ob/ob mice, a distinct model of NAFLD that features microvesicular steatosis and inflammation (fig. S7A). Using RNA-FISH (RNAscope) in GAN animals, we found that *Zfp469* mRNA colocalized with *Col1a1* and *Acta2* transcripts, indicative of aHSCs (fig. S7B) and in agreement with the human MERSCOPE data (Fig. 6D). We broadened this analysis to include other public cohorts of mouse data and confirmed that *Zfp469* mRNA correlated with *Col1a1* expression, and its expression increased upon disease progression in carbon tetrachloride-(CCl_4_-) induced liver disease and bleomycin-induced lung disease (fig. S7C and S7D). Taken together, ZNF469 is a disease-relevant, evolutionarily conserved and generalizable regulator of collagens.

## DISCUSSION

The transition to fibrosis represents a clinically important inflection point in NAFLD morbidity and mortality in humans. We discovered transcriptional regulators of fibrosis in aHSCs using multiomics on human liver samples and a CRISPR genetic screen in a primary cell model. From 100 profiled TFs we found Zinc Finger 469 (*ZNF469*) as the top hit in regulating collagen type 1 protein. Follow-up mechanistic studies at RNA, chromatin and protein levels demonstrated that ZNF469 functions as a positive transcriptional regulator of collagens and matrix genes; a finding conserved in mouse HSCs and in two murine models of NAFLD/fibrosis. In human NAFLD, liver expression of *ZNF469* was positively correlated with the extent of fibrosis. Collectively, these data demonstrate that ZNF469 is a previously unrecognized transcriptional regulator of fibrogenesis in HSCs and NAFLD.

As a member of the C2H2 zinc finger (ZF) protein family, ZNF469 is predicted to function as a TF (*36*). Furthermore, the protein structure possesses multiple nuclear localization sequences and putative transcriptional activation/repression domains. In our study, five lines of experimental evidence strongly indicate that ZNF469 functions as a TF: 1) it is expressed exclusively in the nucleus; 2) CRISPR deletion of *ZNF469* reduces collagen gene expression; 3) the protein localizes to chromatin at collagen gene loci in a ZF-dependent manner; 4) upon *ZNF469* KO, H3K27ac signals — indicative of active enhancers and promoters — are reduced in gene bodies and at distal regulatory elements 5) overexpression of ZNF469 rescues collagen mRNA expression in ZNF469-depleted mouse and human cell lines.

Notably, point mutations in *ZNF469* cause Brittle Cornea Syndrome (BCS) and Ehlers-Danlos syndrome (EDS) in humans (*37, 38*). Both are developmental disorders that feature organ dysfunction from loss of collagen in the eye and musculoskeletal system, respectively. Moreover, deletion of ZNF469 in developing zebrafish decreases synthesis of collagen and proteoglycans (*39*); and knockout of *Zfp469* in mice recapitulates BCS with decreased collagen deposition in the cornea (*30*). No liver phenotypes in the KO mice were described and no adverse effects on viability or fertility were observed (*30*). In addition, it was shown that targeting ZNF469 with siRNAs decreases COL1A1 and COL1A2 in dermal fibroblasts (*40*). These data indicate substantial evolutionary conservation of ZNF469 function in the transcriptional regulation of collagen during development.

Two other TFs prioritized by the CRISPR screen and the multiomics validation data were *RUNX1* and *TBX3*. A recent study revealed that RUNX1 plays an important role in HSC activation and promotes liver fibrosis in a CCl_4_ mouse model by activating TGF-β/SMAD2/3 signaling directly (*41*). Our study confirms that RUNX1 has a profibrotic role in primary human HSCs. Similarly, we also found that TBX3 may be a regulatory factor in HSCs, a finding that could expand its role beyond hepatocytes in which *Tbx3* KO was recently found to protect against high fat diet-induced NAFLD in mice (*42*). In our study, we inferred TF activity using multiple computational tools with paired RNAseq and enhancer profiles as input datasets. Other groups have leveraged global and single-cell gene expression profiles and circulating proteomics in human NAFLD to gain insight into pathways involved in fibrosis (*43, 44*). Across these studies, transcriptomic data consistently stratifies samples by disease stage. Pathways of inflammation, lipid metabolism and ECM are all highly represented. Notably, pair-wise analysis across disease stage groups in one study also enabled identification of a “gene signature” that includes COL1A1 and COL1A2 (*43*)— both regulated by ZNF469. However, ZNF469 was not specifically described in these studies. All of these datasets support the concept that defined molecular mediators likely control key transitions during liver disease progression. Our study adds to this growing body of work by illustrating the computational power of using multiple tools analyzing paired RNA-seq/enhancer datasets to identify regulatory TFs in human NAFLD fibrosis.

In addition to promoter and enhancer elements, ZNF469 bound selectively to collagen gene bodies, where H3K27ac signal was also most significantly lost upon *ZNF469* KO. To our knowledge only PRDM5, in which mutations also cause BCS, has been shown to have such a specific genome occupancy profile at collagens genes (*45*)(*40*). The mechanisms to explain how ZNF469 binds collagen genes remains unclear. No consensus DNA motif has yet been described for this TF in the literature and the large regions of DNA occupancy at gene bodies in our datasets made it more challenging to identify discrete DNA binding sequences. Whether pathogenic variants in ZNF469 causing BCS impact DNA binding directly will be interesting to test as a potential transcriptional mechanism for the collagen downregulation observed in the disease. TFs (e.g. Yin Yang 1, Krüppel-like Factor 4, MYC and GATA2) can bind to RNA through arginine-rich domains (*46*). Thus, structural features of target genes at the RNA level could be determinants of ZNF469 localization. Previous work has also shown that PRDM5 binds to chromatin at collagen gene bodies through direct interactions with RNA polymerase II (Pol II) (*45, 47*). As such PRDM5 and ZNF469 may functionally interact at these gene loci to control Pol II-dependent transcription of collagen.

Our study has limitations. Biopsy samples were taken from individuals undergoing gastric bypass surgery. Consequently, all patients have some degree of systemic metabolic dysfunction associated with obesity including those with normal histology. Furthermore, the demographics, in particular sex and race, are not equally distributed across groups, thereby missing genetic variability across different ethnic groups. A limitation of our CRISPR screen was the focus on TFs. Another group recently performed a genome-wide CRISPR screen to identify TGFβ-responsive mediators of HSC activation, but did not report any results related to *ZNF469* KO(*48*). However, in our screen, an important filtering step was inclusion of TFs identified through analysis of RNA-seq and enhancer landscapes derived directly from human liver samples. Thus, we believe the hits in our screen do have immediate translational relevance to human disease.

Conceptually, TFs are appealing therapeutic targets because of the potential to alter expression and activity of multiple genes in a disease pathway simultaneously in NAFLD. However, TFs are not readily accessible for drug development due to lack of small-molecule binding pockets and presence of intrinsically disordered functional domains (*49*). Nuclear receptor (NRs: PPARs, FXRs, LXRs) are a notable exception. Results of drug studies targeting NRs have been mixed. However, a clinical trial recently demonstrated that a thyroid receptor β agonist can reduce fibrosis score (*4*). Our results indicate that targeted loss-of-function/inhibition of ZNF469 in HSCs should improve liver fibrosis and future studies using preclinical models will directly test this hypothesis. Overall, our work does illuminate the power of a reverse translation approach to discover a previously unrecognized transcriptional mediator governing stage-specific progression of human NAFLD.

## MATERIALS AND METHODS

### Study Design

This study was designed to map and elucidate altered transcriptional circuitries and transcription factor pathways that define the disease states and transitions from healthy to NAFLD and ultimately fibrosis. The outcome of this study are transcription factors that are relevant in long term disease progression accompanied by a deepened understanding of transcriptional landscape and regulation in chronic progressive liver disease at clinically relevant time points.

### Patients and sample collection

Wedge liver biopsies of the left lateral lobe of the liver were obtained at the time of elective bariatric surgery in both male and female patients. Subjects gave informed written consent before participating in this study, which was approved by the Internal Review Board of Vanderbilt University (090657 and 171845) and registered at ClinicalTrials.gov (NCT00983463 and NCT03407833). All studies were conducted in accordance with NIH and institutional guidelines for human subject research. The study protocol conformed to the ethical guidelines of the 1975 Declaration of Helsinki, as reflected in *a priori* approval by Vanderbilt University Medical Center. Liver histology was reported by a minimum of two pathologists with experience in reporting hepatic histopathology. Hepatic steatosis, ballooning, lobular inflammation and fibrosis were reported in accordance with the NASH Clinical Research Network (NASH-CRN) scoring criteria (*50*). Definite histological NASH was defined as NAFLD activity score (NAS) ≥ 5 while borderline NASH was defined as NAS of 3-4. Significant and advanced fibrosis were defined as ≥F2 and ≥F3, respectively. Inclusion criteria included scheduling for bariatric surgery, obesity ≥ 40 kg/m^2^ or ≥ 35 kg/m^2^ and one comorbidity (type 2 diabetes [fasting blood glucose ≥ 120 mg/dL; HbA1C ≥ 6.5%]; known fatty liver disease, hypertension, cardiovascular disease or hyperlipidemia. Patients were excluded if they had prior bariatric surgery, malignancy (< 5 years), known history of intestinal disease, malabsorptive syndrome, were pregnant or breastfeeding, established organ dysfunction, renal disease, alpha 1 ant-trypsin disease, Wilson’s disease, viral hepatis, alcoholic liver disease, or smoked > 7 cigarettes per day.

### Cell culture

All cells were cultured at 37°C in 5% CO_2_ in humidified incubators and were free from mycoplasma (MycoAlert Detection Kit, Lonza). Human primary hepatic stellate cells used for the main experiments were from Lonza (HUCLS, lot HSC190131, female, Caucasian, age = 38 years; BMI = 31.2 kg/m^2^). A second donor was used for confirmation experiments (BioIVT, batch no: S00354, Lot: OTL, male, Caucasian, age = 49, BMI = 26.7 kg/m^2^). Cells were thawed according to the supplier’s instructions and expanded in collagen-coated flasks in human stellate cell growth medium (Lonza, MCST250) supplemented with 50 μM 2-Phospho-L-ascorbic acid (Merck, 49752). Frozen stocks were prepared at passage 4 and used for all subsequent experiments. Typically, cells were thawed and passaged once before electroporation at passage 6. For CRISPR validation experiments cells were seeded in triplicates directly after electroporation, were split at day 7 or 8 into a bigger format, medium was changed at day 4 and day 11 and cells were collected at day 12. For RNAseq experiments, cells were seeded in triplicates in collagen-coated 24 well plates (5000 cells / well) and split into the well of a collagen-coated 6 well plate at day 7. For profiling experiments, cells were first seeded in triplicates in collagen-coated 10 cm dishes after electroporation. To compensate for the differences in cell growth effect of each target, the number of cells plated / 10 cm dish were 166,000 for RUNX1 and TBX3; 100,000 for hNTO, ZNF469 and COL1A1. Each of the 10 cm dish replicates was counted and re-seeded into a collagen-coated T150 flask at day 8 (800,000 cells / flask for RUNX1 and TBX3, 600,000 cells / flask for hNTO, ZNF469 and COL1A1. LX-2 cells were cultured in DMEM (Gibco 11965092) supplemented with 1% penicillin/streptomycin (Gibco 15070063), 50 μM 2-Phospho-L-ascorbic acid (Merck, 49752) and 2% TET-system-approved FCS (Gibco A4736201). JS1 cells were maintained in high glucose DMEM (Gibco 11965-092) supplemented with 2% FBS and penicillin/streptomycin.

### Generation of stable cell lines with inducible ZNF469 constructs

For the generation of stable transgenic LX-2 cell lines, 500,000 cells per well were seeded in a 6-well plate and were transfected with Lipofectamine 3000 according to the manufacturer’s protocol (Invitrogen L3000015). For the transfections, 2.1 μg of the plasmid containing the inducible ZNF469 constructs were combined with 400 ng of the PiggyBac helper plasmid (*51*). Transfected cells were left for 3 days, and subsequently selected with 250 μg/mL Geneticin (Gibco 10131035) for 2 weeks. Medium and Geneticin were replenished every 2-3 days.

### siRNA knock down

JS1 mouse stellate cell line was transfected with a pool of 4 siRNAs targeting Zfp469 (ON-TARGETplus Mouse Zfp469 siRNA, Horizon Discovery, catalogue #L-061504-01-0005) using electroporation (program #8) on the NEON transfection system. A negative control was also included ON-TARGETplus Non-targeting Control Pool (Horizon Discovery catalogue # D-001810-10-05). After electroporation, cells were plated at 10,000 cells per well in a 96-well plate. RNA was extracted for one-step RTqPCR gene expression analysis 24 h after plating.

### Inducible ZNF469 constructs

Human full length ZNF469 coding sequences as well as a version carrying a deletion was cloned into a doxycycline inducible PiggyBac expression vector described previously (*51*). The sequences of the corresponding constructs piHA-ZNF469_FL_PB and piHA-ZNF469_dZF_PB were submitted to GenBank (submission IDs: PP266319 and PP266320, respectively).

### Molecular biology design and analysis software

Geneious Prime was used for construct design and protein sequence analysis. PSORT was used for identification of nuclear localization signals (<u>PSORT: Protein Subcellular LocalizationPrediction Tool (genscript.com))</u>

### RNA extraction and quantification

RNA from human cell cultures was extracted from 6 well plates using RNeasy Plus micro kit (Qiagen, 74034). The plates were washed once with PBS, 350 μl RLT Plus buffer was added to each well and the plates were kept at –80 °C until extraction. Qiagen’s standard protocol including a DNase treatment was followed and the total RNA was finally eluted into 20 μl RNase-free water. RNA quality was assessed using a Tapestation (Agilent) and quantity measured using the Qubit RNA BR assay (Thermo Fisher Scientific, Q10210). For qPCR, 2 μl RNA were used in each PCR reaction using KAPA SYBR FAST One kit (KK4651) and run on a QuantStudio 7 Flex System (Applied biosystems).

Total RNA from frozen liver tissue (30 mg) was isolated using the Qiagen RNeasy Mini kit (Germantown, MD), per manufacturer’s instructions. RNA quality was assessed using the 2100 Bioanalyzer (Agilent). For JS1 cells, RNA was extracted using 40 microliters of Bio-Rad RNA sample preparation reagent (Catalogue #1708898) and frozen for subsequent qPCR with BioRad one-step SYBR green RT-qPCR reagent (Catalogue #1725150). One μl of RNA was used in each PCR reaction.

### RNA-sequencing

NGS libraries were prepared with the Illumina TruSeq Stranded mRNA Sample Preparation kit (Illumina, #20020595) according to the manufacturer’s instructions (Illumina TruSeq Stranded mRNA Reference Guide, # 1000000040498 v00, October 2017), using 400 ng of total RNA as input and applying 12 cycles of PCR at the final enrichment step. The produced NGS libraries were quality controlled using the D1000 ScreenTape and D1000 reagents (Agilent, #5067-5582 and 5067-5583) on the Agilent 2200 TapeStation system. Quantification of the libraries was performed using the Qubit dsDNA HS Assay Kit (Thermo Fisher, #Q32854) on the Biotek FLx800 Fluorescence plate reader. Paired-End 51 bp data were produced using the Illumina NovaSeq 6000 system, generating 26 million reads per sample on average.

The sequencing data were base called with Illumina’s Real-Time Analysis (RTA v3.4.4) software and processed into FASTQ file format, containing the sequence data, with the bcl2fastq2-v2.20.0.422 tool.

### RNA-sequencing analysis

We used the exon quantification pipeline (version 2.5, (*52*)) with STAR (version 2.7.3a, (*53*)) to align the reads against the human genome reference from Ensembl version 98 (*54*) and used PISCES (GitHub - Novartis/pisces) to quantify gene expression.

For liver samples data principal component analysis was performed on the top 5% genes with the highest median absolute deviation using the prcomp R function. Differential gene expression between disease stages (NOR vs NASH_F2/3) was determined using DESeq2 (v1.38.3) with sex as covariate. Genes were defined to be differentially expressed if FDR <= 0.01 and |log2FC| > 0.5. All differential gene-based enrichment analysis was performed with the enricher function from ClusterProfiler (v4.6.2) R package. Cell type enrichment analysis for these differential disease genes was performed with marker gene sets from Aizarani et al (*22*). The HSC gene set was defined by the overlap between NASH_F2/3 genes and the enriched HSC markers from Aizarani et al. Gene ontology analysis for the same differential genes was based on the Hallmark, Wikipathways, Reactome and NABA gene sets from MSigDB (v7.5.1).

For HSCs with CRISPR perturbation differential gene expression and gene set enrichment analysis was performed as described in section above. Here we compared CRISPR knockouts with non-targeting control (hNTO). For the gene set enrichment analysis we used the gene ontology biological processes gene sets from MSigDB (v7.5.1).

The CRISPR editing analysis was performed by first matching back the location of gRNAs to the human genome using the vmatchPattern function from the R-package Biostrings (v2.64.1). Genotyping of the gRNA locations was performed using the pileup function from Rsamtools (v2.14.0) with following parameters: distinguish_strands = FALSE, distinguish_nucleotides = TRUE, ignore_query_Ns = TRUE, include_deletions = TRUE, include_insertions = TRUE.

### CUT&RUN chromatin profiling

CUT&RUN was carried out as described previously with the following modifications (*55*). For CUT&RUN on liver tissues flash-frozen pieces of human liver biopsies were weighted (around 1 g per sample) and transferred to pre-cooled tissue TUBES TT1 (Covaris, PN 520128). Tissues were pulverized with the CP02 cryoPREP automated Dry Pulverizer from Covaris by executing one pulse at intensity level 2 followed immediately by a second pulse at intensity level 4. While constantly kept on dry ice, around 5 mg of tissue powder per donor were transferred into BSA-pre-coated and pre-cooled 1.5 mL Eppendorf tubes. Tissues were washed with 1 mL of cold DPBS and centrifuged for 5 min at 1000 x g (4 °C). Tissues were fixed for 5 min at room temperature (RT) with 1% fresh formaldehyde (Pierce 16% Formaldehyde Methanol Free, 28906) diluted in PBS under gentle rotation. The fixation was quenched with 1:10 vol of 1.25 M glycine solution for 5 min with gentle rotation. Samples were washed again with DPBS and resuspended in 1 mL of NE Buffer (20 mM HEPES pH 7.5, 10 mM KCl, 0.5 mM Spermidine, 0.1 % Triton X-100, 20% Glycerol, cOmplete EDTA-free Protease Inhibitor Cocktail, 4693132001). Samples were transferred into Lysing Matrix D 2 mL tubes containing 1.4 mm ceramic beads (MP Biomedicals, 116913050-CF) and treated for 10 sec at 5000 rpm with the Precellys 24 Tissue Homogenizer from Bertin Technologies. Cell suspensions were quickly placed on ice for 5 min and filtered through a 70 µm cell strainer, then transferred into new BSA-pre-coated 1.5 mL Eppendorf tubes and centrifuged for 3 min at 500 x g. Finally, pellets were resuspended in 100 µL of Wash 1 Buffer (20 mM HEPES pH 7.5, 150 mM NaCl, 0.5 mM Spermidine, cOmplete EDTA-free Protease Inhibitor Cocktail, (4693132001) and further processed as described below.

For cell cultures cells were detached with 0.25% Trypsin-EDTA (Gibco, 25200056) for 5 min at 37°C, centrifuged at 300 x g for 3 min, washed once with PBS and counted with Countess cell counter (Thermo Fisher). The fixation with formaldehyde and the quenching were done as described above. Cells were washed with PBS twice, then re-suspended in PBS and transferred into a final volume of 500 μL in 1.5 mL Eppendorf tubes pre-coated overnight with 1% BSA. Aliquots of 2 million-, or 200,000-cells were used for the ZNF469, or for the H3K27ac profiling, respectively, and processed as described below.

For all sample sources processed as described above, Activated BioMag® Plus Concanavalin A magnetic beads (Bangs Laboratories, BP531) were added to the cell suspension (40 µL for ZNF469 or 10 µL for H3K27ac) and incubated for 10 min with gentle rotation. After magnetic separation, cells were resuspended in 200 µL of DIG**-**wash EDTA Buffer (20 mM HEPES pH 7.5, 150 mM NaCl, 0.5 mM Spermidine, 0.02% Digitonin, 2 mM EDTA, cOmplete EDTA-free Protease Inhibitor Cocktail) with the primary antibody for 2 h with gentle rotation. Primary antibodies used were rabbit polyclonal ZNF469 (Sigma, HPA069784), polyclonal H3K27ac (Abcam, ab4729). Cells were washed twice with 1 mL of DIG-wash EDTA buffer and incubated for 1 h at 4 °C with gentle rotation in 200 µL of DIG-wash EDTA Buffer with CUTANA pAG-MNase (protein A-protein G-Micrococcal Nuclease fusion, Epicypher, 15-1016). Subsequently, the cells were washed twice with 1 mL of DIG-wash EDTA Buffer and once with 1 mL of DIG-wash Buffer without EDTA. For the pAG-MNase digestion, the cells were re-suspended in 100 µL of cold DIG-wash Buffer without EDTA, then CaCl_2_ was added to a final concentration of 2 mM to activate the MNase. Samples were incubated for 30 min at 0°C in an aluminum block. Digestion was terminated by adding 100 µL of 2X Stop Buffer (400 mM NaCl, 20 mM EDTA, 4 mM EGTA, 0.02% Digitonin, 100 µg/mL RNase A, 50 µg/mL Glycogen) to the samples. Samples were incubated for 30 min at 37 °C without shaking. Supernatants were collected and treated with SDS (1% final concentration) and 200 µg/mL Proteinase K for minimum 4 h at 65°C. DNA was extracted by phenol-chloroform for the ZNF469 profiling to keep the small DNA fragments and with the DNA Clean & Concentrator-5 kit (Zymo Research, D4004) for the H3K27ac profiling. Purified DNA was eluted with the EB buffer (Qiagen, 19086) and transferred in 1.5 mL DNA LoBind tubes (Eppendorf, 30108051).

For library preparation, the NEBNext Ultra II DNA Library Prep kit for Illumina (New England Biolabs, E7645L) was used as described previously (*56, 57*). The end repair step was adapted for each target: 30 min at 20°C and then 1 h at 50°C for ZNF469; 30 min at 20°C and then 30 min at 65°C for H3K27ac. The NEB adaptor was diluted at 1:15. The NEBNext Multiplex Oligos for Illumina SET2 (New England Biolabs, E7500L) were used for the indexing. The PCR reaction was adapted for each target: 9 cycles and annealing / extension for 10 sec at 65°C for ZNF469; 8 cycles and annealing / extension for 75 sec at 65 °C for H3K27ac. Libraries were purified using Agencourt AMPure XP beads (Beckman Coulter, A63881) and the concentrations and profiles were checked on TapeStation with a D1000 Screen Tape. Libraries were sequenced using NovaSeq6000 paired-end mode 2 x 51 bp.

### CUT&RUN data analysis

For the H3K27ac liver cohort principal component analysis was performed on the top 5% peaks with the highest median absolute deviation using the prcomp R function. Like RNA-seq, differential H3K27ac signal between NOR and NASH_F2/3 was tested using DESeq2 (v1.38.3) with sex as covariate. We defined differential H3K27ac regions as FDR <= 0.01 and |log2FC| > 0.5. Metaprofiles and IGV tracks of CUT&RUN and other chromatin signals were computed using a custom script which is based on the ScoreMatrix function from genomation (v1.19.1) R-package. Cell type specific scATAC-seq peaks were obtained from Zhang et al (*58*). NOR and NASH specific differential regions overlapped with these peaks. Resulting confusion matrices were used to test for enrichment with two-sided Fisher’s exact tests.

For H3K27ac in HSCs, differential H3K27ac between hNTO and ZNF469 KO was determined as described in the previous section. Peaks were identified as differentially acylated if FDR <= 0.01 and |log2FC| > 0.5. HSC ZNF469 KO RNA-seq and H3K27ac CUT&RUN were integrated by comparing the log2FC of gene expression and H3K27ac at the promoter of a gene or distal intergenic peaks were assigned to genes using the capture Micro-C interactions.

For ZNF469 in HSCs, ZNF469 binding was determined against a knockout control using DESeq2 (v1.38.3) to compare ZNF469 CUT&RUN hNTO against ZNF469 KO. Since we expect genome-wide changes in TF binding, we computed size factors based on *E. coli* spike in reads as previously described. ZNF469 binding sites were defined using following thresholds FDR <= 0.05 and |log2FC| > 0.5.

ChIPseeker (v1.34.1) was used to annotate the binding sites with their genomic locations. Enrichment of the ZNF469 binding sites was determined by comparing their genomic location with the ones of non-significant bound sites using a one-sided Fisher’s exact test.

### Micro-C and promoter capture

To generate a genomic promoter interaction map in human primary hepatic stellate cells (HSC190131) two replicas of 1-1 million cells were fixed with disuccinimidyl-glutarate and processed with the Dovetail Micro-C Kit (21006) following a promoter capture step (Dovetail Human Pan Promoter Enrichment Kit, 25013) according to the manufacturer’s manual. For two replicates 494 million fragments were sequenced on the Illumina NovaSeq 6000 system. The above reagents as well as Dovetail Library Module for Illumina (25004), Dovetail Dual Index Primer Set #1 for Illumina (25010), and the Promoter Panel Informatics Analysis service (8015) were from Dovetail Genomics.

### Micro-C and promoter capture analysis

Capture Micro-C preprocessing was performed as outlined at https://dovetail-capture.readthedocs.io/. Briefly, paired-end reads were aligned to hg38 using bwa meme (v2.2.1) with following parameters: −5SP-T0. Resulting alignment was searched for valid ligation events using the parse function from the pairtools pipeline (v1.0.2) with following parameters: --min-mapq 40 –walks-policy 5unique –maxinter-align-gap 30. PCR duplicates were removed using pairtools dedup and bam files for downstream analysis were generated using samtools (v1.13). Chromatin-chromatin interactions calling was performed using CHiCAGO (v1.26.0). Therefore, bait fragment (BF) and other end fragments (OEF) maps were generated at a 5kb resolution. These maps were used to compute the coverage from the bam files using the bam2chicago.sh script. Resulting compatible input files were used for interaction calling with CHiCAGO. Interactions with a score higher than 5 were considered significant. Bait plots were plotted based on CHiCAGO generated statistics with custom ggplot2 (v3.4.4) based R code.

### Analysis to identify transcription factor candidates

To identify candidate TFs different methods were used with the appropriate input data as described below.

#### ANANSE (*23*)

First, we identified nucleosome free regions (NFRs) per patient using HisTrader (github.com/SvenBaileyLab/HisTrader) with default parameters and H3K27ac CUT&RUN data as input. We combined these NFRs per disease stage (NOR, NASH_F2/3) by merging the bed files using BEDtools (v2.27.1). TF binding to disease stage NFRs were predicted using ANANSE binding (default settings) in combination with the merged locations and H3K27ac CUT&RUN signals. Next, gene regulatory networks per disease stage were reconstructed using these TF binding predictions in combination with gene expression measurements from RNA-seq as input for the ANANSE network function. Finally, TFs were prioritized using ANANSE influence (default settings) with the DESeq2 differential genes between NOR and NASH_F2/3, the gene regulatory networks (GRNs) for NOR AND NASH as input.

#### CRCmapper (*24*)

We identified super-enhancers from H3K27ac CUT&RUN for each patient using ROSE2 (bitbucket.org/young_computation/rose) with default settings. These were used to construct Core Regulatory Circuitries (CRCs) using CRCmapper (github.com/linlabcode/CRC) in combination with NFRs as subpeaks (see ANANSE) and actively expressed genes as defined by matched RNA-seq (TPM > 1). Resulting patient level CRCs were used as input to compute various network statistics (in degree, out degree, total degree, betweenness, alpha centrality and eigenvector) using the R-package igraph (v1.3.4). We compared the out degree network statistics between NOR and NASH_F2/3 using Wilcoxon rank sum tests.

#### DoRothEA (v1.10) (*25*)

Curated GRNs for Homo sapiens were queried via the DoRothEA R-package. To compute a TF activity score per patient from RNA-seq, we used these GRNs in combination with the TPM matrix as input for the run_ulm function from DecouplR (v2.4; minsize = 5). Resulting TF activity scores per patient were summarized by comparing NOR vs NASH_F2/3 using Wilcoxon rank sum tests.

#### Homer (v4.11) (*28*)

The motif consensus library from Lambert et al was used for the motif enrichment with HOMER. The analysis itself was performed using the function findMotifsGenome with NASH_F2/3 as foreground and NOR differential regulatory region as background sequences.

#### MonaLisa (v1.2) (*27*)

Motif enrichment with MonaLisa was performed using the motif consensus library from Lambert et al. Motif counts were obtained by matching the Lamber et al PWMs to the H3K27ac consensus peaks using the matchMotif function from motifmatchr. We computed GC and CpG observed/expected ratios were computed as additional features. The motif counts and ratios were used to model the H3K27ac log2(FC) between NOR and NASH_F2/3 using the randLassoStabSel function from MonaLisa (cutoff = 0.8).

#### RcisTarget (v1,18.2) (*26*)

We used RcisTarget to compute the motif enrichment for NASH_F2/3 differential regions. Coordinates of differential regions were first converted from hg38 to hg19 using liftOver (v1.58). Region based motif enrichment was performed using the hg19 ranking database and NOR peaks as background. The analysis was performed by running the cisTarget function from RcisTarget.

### sgRNA stocks preparation

For the arrayed CRISPR screen, crRNAs were either cherrypicked from Horizon’s Human Edit-R Drug targets (GC-004650-05) or Human Druggable Subset (GC-004670-05) synthetic crRNA libraries or were purchased from IDT. Individual crRNAs from the Horizon libraries (0.5nmol/well) were resuspended into 35 μl 10mM Tris pH7,4 to generate a 15 μM solution using a Multidrop Combi reagent dispenser (ThermoFisher Scientific), placed on an orbital shaker for a few minutes and incubated at RT for about 1h. They were cherrypicked from the original library plates and 4 crRNAs per target were pooled (4 x10 μl) and re-arrayed using a Biomek-FX automated pipetting system (Beckman Coulter). Edit-R CRISPR-Cas9 Synthetic tracrRNA (Horizon Discovery Biosciences, U-002005-20) was resuspended and diluted to 15 μM in 10 mM Tris pH 7.4 and 40 μl were then added to the crRNAs using a Multidrop Combi dispenser and incubated at RT for 20 min. The guides were then transferred to stock plates using a CyBio SELMA 96/60ul semi-automated pipette (Analytik Jena) according to the final layout and kept at −20°C (final concentration of 7.5 μM). For all targets that were not available from the Horizon libraries, crRNAs were purchased from IDT as a pool of 4 x 2 nmol crRNAs (Alt-R CRISPR-Cas9 crRNA, sequences according to T.spiezzo library, (*59*)). They were resuspended in 10mM Tris pH 7.4 to obtain 100 μM stocks, further diluted to 15 μM and incubated for 20min at RT with 15 μM Edit-R CRISPR-Cas9 Synthetic tracrRNA. They were finally transferred to the stock plates using a multichannel pipette according to the final layout and kept at −20°C. For simplicity the complexed crRNA-tracrRNA molecules will be referred to as “sgRNA” in the manuscript.

### RNP preparation and delivery

For the arrayed CRISPR screen, 7.5 μM sgRNA stock plates were thawed and 5.76 μl of each guide was transferred to a 96 well conical bottom plate (Nunc, 249935) using a CyBio SELMA. Control sgRNAs were added manually. Alt-R S.p. Cas9 Nuclease V3 (IDT, 1081059) was diluted to 15 uM in 10 mM Tris-HCl pH 7.4 and 2.4 μl were dispensed on top of the sgRNAs using a Mantis microfluidic liquid dispenser (Formulatrix). The RNP complexes were incubated at RT for 15min and then kept at 4°C until use. During this time, human HSCs were detached and resuspended in P3 buffer (Lonza, P3 primary cell 384-well nucleofector kit) containing Alt-R Cas9 Electroporation Enhancer (IDT, 1075916). 15.84 μl of cells were then pipetted onto the prepared RNP complexes using a multichannel pipet (36 pmol RNP for 30,000 cells) and 20 μl of that mix was transferred to a Lonza 384-well nucleofector plate (30 pmol RNP and enhancer for 25,000 cells per condition) and electroporated using program CA-137 on a 384-well Nucleofector system.

40 μl fresh medium were added to the electroporation plate and 2 μl cells were transferred into prefilled and prewarmed collagen-coated 96 well plates (Biocoat Collagen I 96-well clear flat bottom TC-treated microplates, Corning, 356698) in multiple replicates using a Cybio SELMA (final seeding density of 800 cells/well). The plates were incubated at 37°C and a medium change was performed 4 days after electroporation. The plates were then processed as detailed in the other sections.

Conditions were adapted and scaled up for each of the validation experiments according to the desired final cell amounts. RNP ratios per cell were kept constant and ranged from 30 pmol RNP for 25,000 cells to 480 pmol RNP for 400,000 cells. Seeding densities were also scaled up to mimic the conditions of the screen and fresh crRNAs were purchased from IDT for experiments requiring more concentrated RNPs.

For CRISPR targeting of *Zfp469* in JS1 cells, RNPs were generated using two crRNAs per region of the gene with one gRNA pair targeting the proximal region and another pair targeting the zinc finger region (base pairs 9538-9661) of *Zfp469* open reading frame. 3 μl of crRNA and tracrRNA (stocks 100 μM) were annealed together at 95°C in 4 μl of R buffer (NEON transfection system, Thermo Fisher, Cat. #MK10096). After cooling for 10 mi at RT, 1.5 μl of Cas9 nuclease (spCas9, IDT #1081058, stock 10 mg/mL) was incubated with crRNA/tracrRNA for 30 minat 20°C. During this incubation, cells were collected and counted for electroporation at a concentration of 1.25e6 cells per 110 μl of R buffer. Electroporation was performed per manufacturer protocol (NEON) in 3 mL buffer E2 per tube with 100 μl tips after mixing RNP with cells. Protocol used for electroporation: Pulse Voltage = 1300, Pulse Width = 20, Pulse Number = 2.

### Immunofluorescence staining

All the steps were done at RT. For imaging cells in micro-well plates (Greiner Bio-One 384 Collagen type I CELLCOAT 384 well, 781956) cells were fixed directly by adding paraformaldehyde (PFA) (Electron Microscopy Sciences, 15714S) to the medium (final concentration 4%) for 15 min on a shaking platform. Cells were washed (80 μl) 3 times with PBS by using the Washer Dispenser EL 406 from Biotek. Cells were permeabilized with Blocking Buffer (PBS pH 7.5, 2 % BSA, 0.1% Triton X-100) for 45 min, followed by 3 washes with Wash 2 Buffer (PBS pH 7.5, 0.1% Triton X-100). Cells were then incubated with the primary antibody in Blocking Buffer for a minimum of 3 h. For COL1A1 staining the monoclonal mouse anti human COL1A1 antibody (Developmental Studies Hybridoma Bank, M-38, lot 5/13/2021) was used at a dilution of 1:1500. After 3 washes with Wash 2 Buffer, the cells were incubated for 1 h with the secondary antibody (goat anti-mouse Alexa Fluor 488, Invitrogen, A11029) diluted at 1:1000 in the Blocking Buffer supplemented with Hoechst diluted at 1:10,000. Finally, cells were washed 3 times with the Wash 2 Buffer (PBS pH 7.5, 0.1% Triton X-100) and stored with aluminum sealing in PBS at 4°C until imaging. For LX-2, cells plated in 96-well plates (Greiner 96-well microplates, µclear®, 655097) were fixed in 100 μl PBS + 4% PFA for 15 min at RT and washed 3 times with 150 μl of PBS. All the subsequent steps were done as described above. The anti-ZNF469 antibody (Sigma, A31572, lot R98565) was diluted 1:400. Secondary antibody (donkey anti-rabbit Alexa Fluor 555, Invitrogen, A31572) dilution was 1:1000. DAPI (Sigma D9542, lot 175158) was used at a final concentration of 1 ng/mL.

### Image acquisition, processing and quantification

For the CRISPR screen the immunofluorescent images were captured with the confocal dual spinning disk Cell Voyager CV7000 and quantified with High Content Analysis Software CellPathFinder (Yokogawa). Confocal images of LX-2 cells were captured with ZEN 3.2 on the Zeiss Axio Observer Z1 microscope with 63x magnification (Plan-Apochromat 63x/1.40 Oil DIC M27) and images were processed with Fiji (*60*). Confocal images from RNAscope experiments were collected on a Zeiss LSM880 using a 63x/1.40 Plan-APOCHROMAT OIL objective.

### RNAscope

For RNAscope of mouse livers, mice were perfused with 4% PFA for 10 min and tissues were harvested and put in 4% PFA for 24 h at 4°C. After fixation, samples were paraffin embedded and sectioned (i.e. FFPE). FFPE liver tissues were subsequently stained using the RNAscope Multiplex Fluorescent Detection Reagents_v2 (ACDBio, catalogue #323110). Target retrieval was performed for 30 min at 100°C. Protease treatment used was Protease Plus for 30 min at 40°C. RNAscope probes used were Mm-Col1a1-C3 (ACD #319371-C3), Mm-Acta2 (ACD #319531), Mm-Zfp469-C2 (ACD #1142291-C2, detection by Opal Fluorophores (Akoya Biosciences - Opal 520 #FP1487001KT, Opal 570 #FP1488001KT Opal 690, #FP1497001KT).

### MERSCOPE sample processing

Five micron thick sections of FFPE human liver specimens were prepared and placed in groups of four on round glass slides provided by Vizgen Corp. (MERSCOPE slide part number 20400101). One mL of a fiducial premix dilution (1:500 in 1XPBS) was added on top of each MERSCOPE slide. Slides were dried at 55°C for 15 min then 2 h at RT before overnight shipment on dry ice to European Spatial Biology Center (Leuven, Belgium) for routine processing.

### MERSCOPE analysis

For the cell-segmentation task, we used Vizgen Post-processing Tool (VPT, v1.0.2) with python (v3.9.6). Cell segmentation was performed on DAPI, polyT and Cellbound2 staining from the z1-plane using the “segmentation family” Cellpose (*61*) in cyto2/2D mode with a cell diameter of 100 μM. For the transcript-to-cell assignment, we used maximal projection from the different z-planes to the respective cells determined in layer z1. With VPT we obtained the cell-metadata and the cell-by-gene matrix files. Cells were filtered for at least 5 different detectable transcripts and only samples with more than 20,000 remaining cells were used for downstream analysis.

Transcript x cell matrices from different patients were merged and downstream analysis performed using Seurat (v5.0.1). Briefly, transcripts were normalized using the SCTransform function with the setting clip.range = c(−10,10) followed by dimensionality reduction via PCA. We chose the first 20 PCs to perform the UMAP embedding (RunUMAP; default) and shared nearest-neighbor graph construction (FindNeighbors; default). Cells were clustered using FindClusters with resolution = 0.2. Resulting clusters were manually annotated based on expression of marker genes in our probe panel.

To determine the co-expression of *ZNF469* with all other transcripts, we binarized the transcript expression and performed one-sided Fisher’s exact tests to test the enrichment of other transcripts in *ZNF469* positive cells. This was done on an individual patient level and over the combined analysis of all samples.

## Statistical analysis

Primary data, sequences and key reagents are presented in table S1-S10. Statistical analysis was performed using R (3.4.2) for omics data. For the CRISPR data we used a Wilcoxon test to compare the medians of the COL1A1 immunofluorescence expression. qPCR data (JS1 CRISPR, JS1 siRNA, HSC CRISPR) were analyzed with one-way ANOVA and represented using GraphPad Prism 10.0 software (* = P ≤ 0.05; ** = P ≤ 0.01, *** = P ≤ 0.001, **** = P ≤ 0.0001). Sequence data analysis was performed using R 4.2.1. Additional details of statistical tests used for each experiment are described in the Material and Methods section.

### Data and materials availability

#### This study

This study:
| GEO | Name | Description | Assay | Species |
| --- | --- | --- | --- | --- |
| **Will be updated** | RNAseq in liver cohort | RNAseq in liver cohort | RNA-seq | hs |
| **Will be updated** | CUT&RUN in liver cohort | Chromatin profiling of H3K27ac in liver cohort | CUT&RUN | hs |
| **Will be updated** | RNAseq in HSCs | RNAseq in HSCs after knockout | RNA-seq | hs |
| **Will be updated** | CUT&RUN of H3K27ac in HSCs | Chromatin profiling of H3K27ac in HSCs | CUT&RUN | hs |
| **Will be updated** | CUT&RUN of H3K27ac in HSCs upon ZNF469 KO | Chromatin profiling of H3K27ac in HSCs with ZNF469 knockout | CUT&RUN | hs |
| **Will be updated** | CUT&RUN of ZNF469 in HSCs | Chromatin profiling of ZNF469 | CUT&RUN | hs |
| **Will be updated** | CUT&RUN of ZNF469 overexpression in LX2 | Chromatin profiling of ZNF469 with overexpression constructs | CUT&RUN | hs |
| **Will be updated** | MicroC-promoter capture in HSCs | MicroC-promoter capture uses probes to pull down promoters from the sequencing library | MicroC | hs |
| **Will be updated** | RNAseq CDAHFD mouse model | RNAseq in a choline-deficient, amino acid defined high fat diet (CDAHFD) mouse model | RNA-seq | mm |
| **Will be updated** | RNAseq GAN mouse model | RNAseq in a Gubra Amylin (GAN) mouse model | RNA-seq | mm |
| **Will be updated** | RNAseq ob/ob mouse model | RNAseq in an ethionine-treated ob/ob mouse model | RNA-seq | mm |

#### Public studies

Public studies:
| GEO | description | Assay | Species | Ref |
| --- | --- | --- | --- | --- |
| GSE184462 | Whole human tissues<br>scATAC-seq | scATAC-seq | hs | A single-cell atlas of chromatin accessibility in the human genome - ScienceDirect |
| GSE135251 | Liver NASH patient<br>cohort | RNA-seq | hs | Transcriptomic profiling across the nonalcoholic fatty liver disease spectrum reveals gene signatures for steatohepatitis and fibrosis \| Science Translational Medicine |
| GSE130970 | Liver NASH patient<br>cohort | RNA-seq | hs | Gene Expression Predicts Histological Severity and Reveals Distinct Molecular Profiles of Nonalcoholic Fatty Liver Disease \| Scientific Reports (nature.com) |
| GSE126848 | Liver NASH patient<br>cohort | RNA-seq | hs | https://journals.physiology.org/doi/full/10.1152/ajpgi.00358.2018 |
| GSE150910 | Lung IPF patient<br>cohort | RNA-seq | hs | https://www.atsjournals.org/doi/10.1164/rccm.2020-01-0134OC |
| GSE173355 | Lung IPF patient<br>cohort | RNA-seq | hs | https://www.atsjournals.org/doi/10.1165/rcmb.2020-0546OC |
| GSE134692 | Lung IPF patient<br>cohort | RNA-seq | hs | RNA sequencing of transplant-stage idiopathic pulmonary fibrosis lung reveals unique pathway regulation - PMC (nih.gov) |
| GSE152329 | CCL4 Liver fibrosis<br>mouse model | RNA-seq | mm | https://www.sciencedirect.com/science/article/pii/S2352345X2030134X |
| GSE195773 | IPF Bleomycin mouse<br>model | RNA-seq | mm | Time and phenotype-dependent transcriptome analysis in AAV-TGFβ1 and Bleomycin-induced lung fibrosis models \| Scientific Reports (nature.com) |

## Author contributions

Conceptualization: MF, CRF, BF, JDB, CK

Methodology: SS, FW, NL, NP, DE, TW, AR, IC, PT, JK, MA, MM, LMD, QS, DPB, YX, PM, AR, SB, SW, JL, WJE, BW, JDB, CRF, BF, CK

Investigation: SS, JDB, CRF, BF, CK Visualization: SS, BF, CK

Funding acquisition: MF, JDB, CRF Supervision: MF, CM, DH, FN Writing – original draft: JDB, CK

Writing – review & editing: SS, JDB, CRF, BF, CK

## Acknowledgments and Funding

We thank Jochen Singer, and Florian Kiefer for RNAseq and chromatin pipeline development; Manuel Scheidmann and Ralph Riedl for CRISPR help; Jasmin Hägele and Ulrike Naumann for supporting NGS sequencing; Gabi Schutzius, Adrian Salathe, Matthias Müller, Melanie Pellisson, Linda Greenbaum, Iwona Ksiazek, Jan Tchorz, Frederic Bassilana and Heinz Ruffner for scientific advice; Michael Hoever, Richard Janovjak, Paul Schroeder, Lachlan Campbell, Sarah Bardouille, and John Dotson for operational and legal support; and Ulrich Schopfer and John Tallarico for financial and organizational support. <u>Funding</u>: Core Services performed through Vanderbilt University Medical Center’s Digestive Disease Research Center supported by NIH grant P30DK058404. The Vanderbilt Translational Pathology Shared Resource is supported by NCI/NIH Cancer Center Support Grant 5P30 CA68485-19. The Diabetes Research Training Center is supported by DK020593 (to the Vanderbilt Diabetes Research and Training Center)

## List of Supplementary Materials

**Fig. S1 to S7**

**Table S1 to S10**

**Supplementary Table 1:** Clinical characteristics of the study population

**Supplementary Table 2:** Differential genes for all RNA-seq datasets

**Supplementary Table 3:** Differential peaks for all CUT&RUN datasets

**Supplementary Table 4:** TF tool results

**Supplementary Table 5:** CRISPR guides

**Supplementary Table 6:** CRISPR raw results

**Supplementary Table 7:** CRISPR statistical results

**Supplementary Table 8:** ZNF469 sequence information

**Supplementary Table 9:** MERSCOPE sample information and probes

**Supplementary Table 10:** Key reagents

