## Supplemental Figures for "The transcription factor ZNF469 regulates collagen production in liver fibrosis"

Supp Figure 1: Integration of transcriptomics and cis-regulatory landscapes in human NAFLD livers predicts activity of transcription factors involved in fibrosis

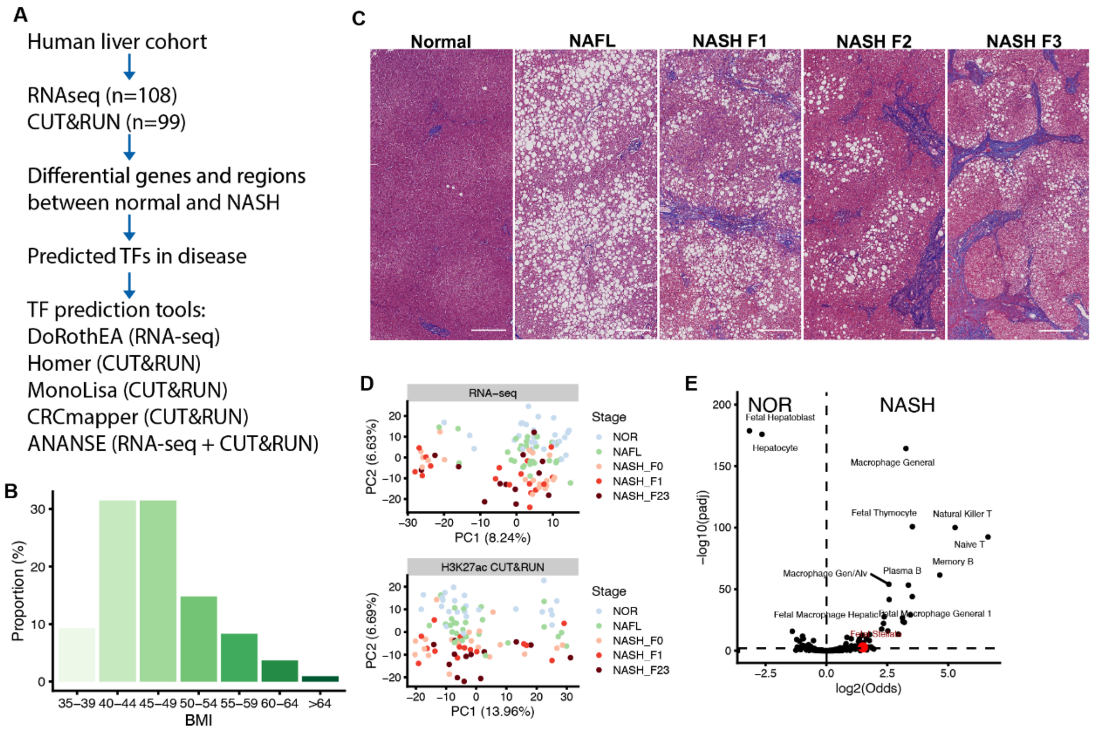


**Supplementary Fig. 1:**

(**A**) Schematic of multiomics data analysis workflow to identify potential transcription factors involved in disease progression. (**B**) Histogram of body-mass index (BMI) for the liver cohort (n=108). (**C**) Representative photomicrographs of Masson's Trichrome stained NASH liver biopsy samples demonstrating histology representative of normal (KUV229), NAFL (ERH825), F1 fibrosis (OTZ834); (2) F2 fibrosis (YIU233) and (3) F3 fibrosis (DNM623). Scale bar = 100 microns. Slides were reviewed and scored by pathologists who were blinded to patient details and clinical status. (**D**) PCA of RNAseq and CUT&RUN colored by histopathology. (**E**) Volcano plot of scATACseq cell type specific peak enrichment in normal vs NASH.

Supp Figure 2: CRISPR loss-of-function screen identifies transcriptional regulators of collagen production in primary human hepatic stellate cells.

**A**

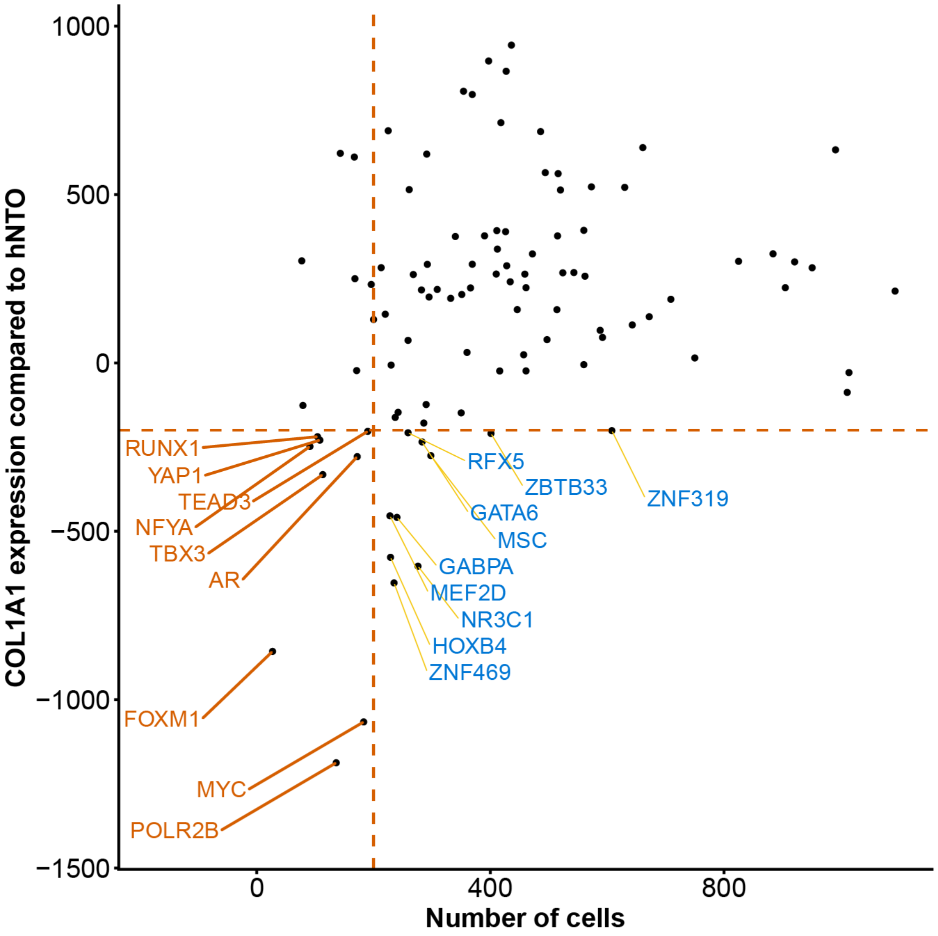


#### Supplementary Fig. 2.

**(A)** Scatter plot showing cell count vs COL1A1 immunofluorescence.

Supp Figure 3: ZNF469 knockout alters collagen mRNA expression in HSCs.

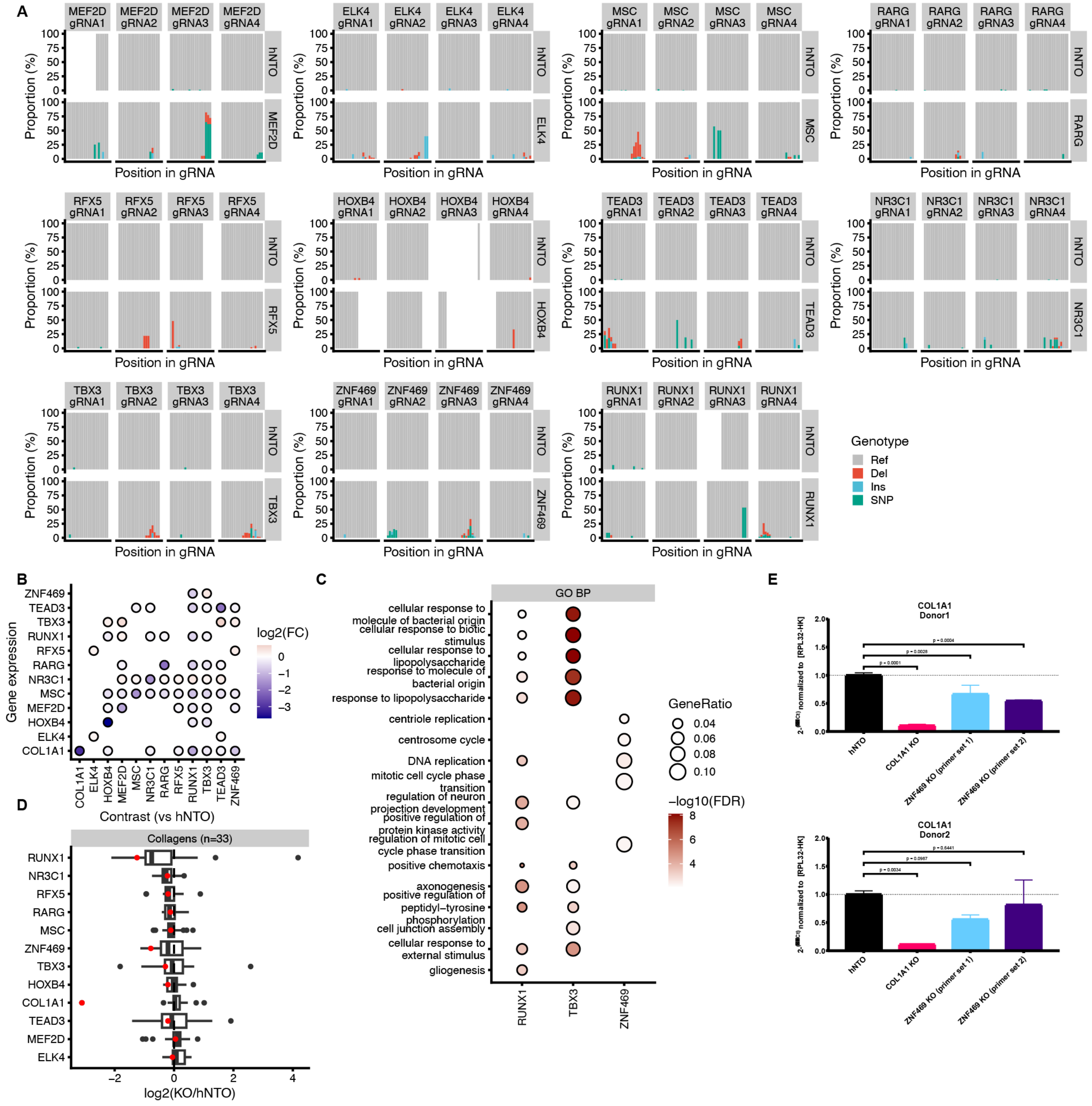


#### Supplementary Fig. 3:

**(A)** RNAseq indel plot shows editing confirmation for *MEF2D, ELK4, MSC, RARG, RFX5, HOXB4, TEAD3, NR3C1, TBX3, ZNF469, RUNX1*. (**B**) Dot plot of TF expression after KO of top hits from the CRISPR Screen. **(C)** Gene set enrichment of upregulated genes upon *ZNF469*, *RUNX1* or *TBX3* KO. **(D)** Gene Set enrichment of upregulated genes upon

*ZNF469*, *RUNX1* or *TBX3* KO. **(E)** Bar plot of COL1A1 expression in 2 separate donors of primary HSCs after ZNF469 CRISPR KO. (see table S5 for sgRNA sequences)

**Supp Figure 4: ZNF469 is a TF.**

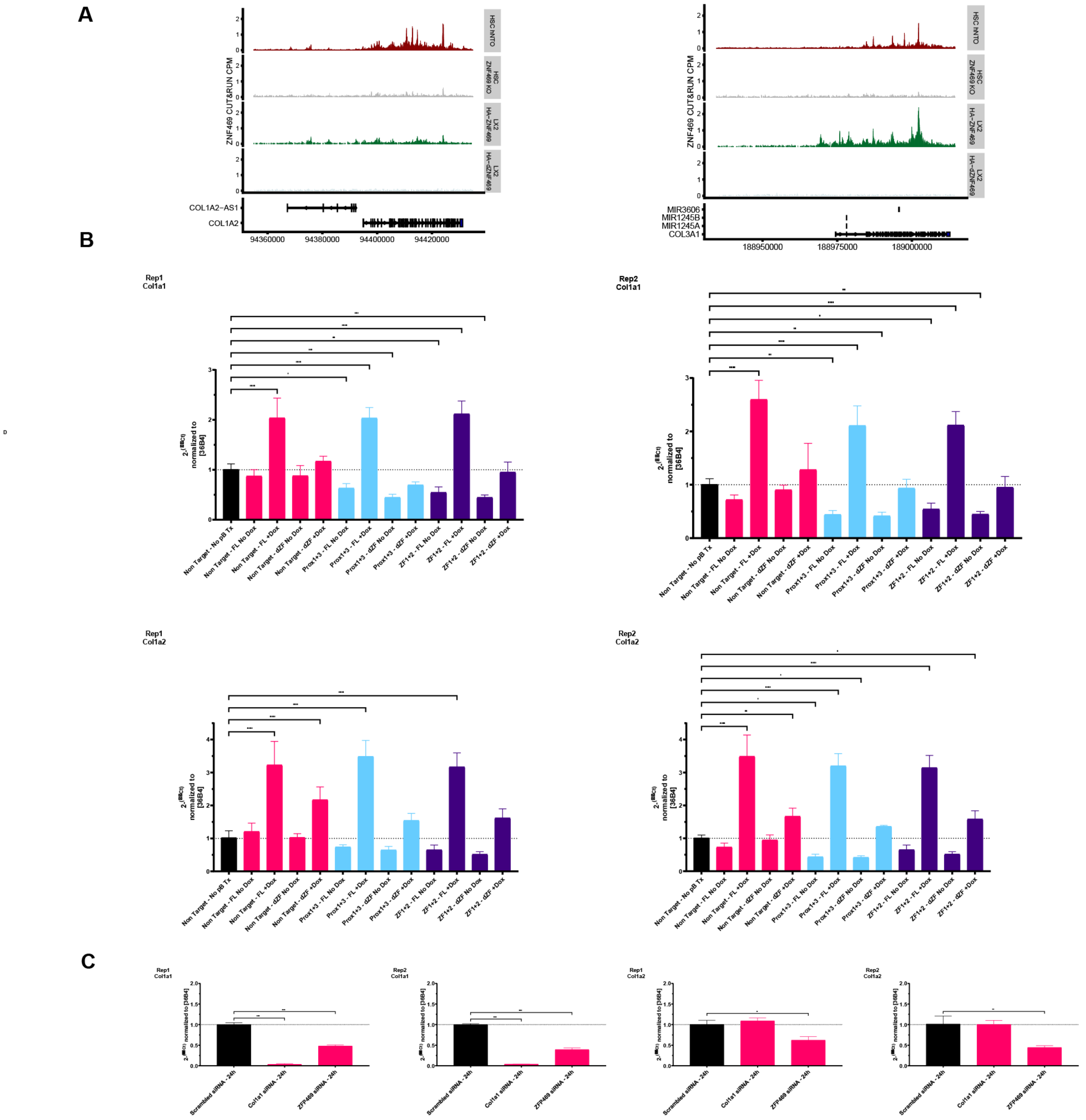


### Supplementary Fig 4:

(**A**) Genome browser tracks at the *COL1A2* and *COL3A1* locus of ZNF469 CUT&RUN signals generated with the anti-ZNF469 antibody in CRISPR experiments in none-targeting and ZNF469 targeted human hepatic stellate cells and transgenic LX-2 cells with doxycycline inducible full length or deletion harboring ZNF469 cDNA (**B**) Bar plots of *Col1a1* and *Col1a2* mRNA expression 7 days after *Zfp469* CRISPR KO in mouse JS1 cells and ZFP469 rescue by overexpression, 24 h after induction of transgene with doxycycline. One gRNA pair targeting the proximal region and another pair targeting the zinc finger region (base pairs 9538-9661) of *Zfp469* open reading frame. (**C)** Bar plots of *Col1a1* and *Col1a2* reduction following *Zfp469* knockdown with siRNA (24 h) in JS1 cells.

**Supp Figure 5: ZNF469 knockout alters local chromatin structure at collagen and ECM loci in HSCs.**

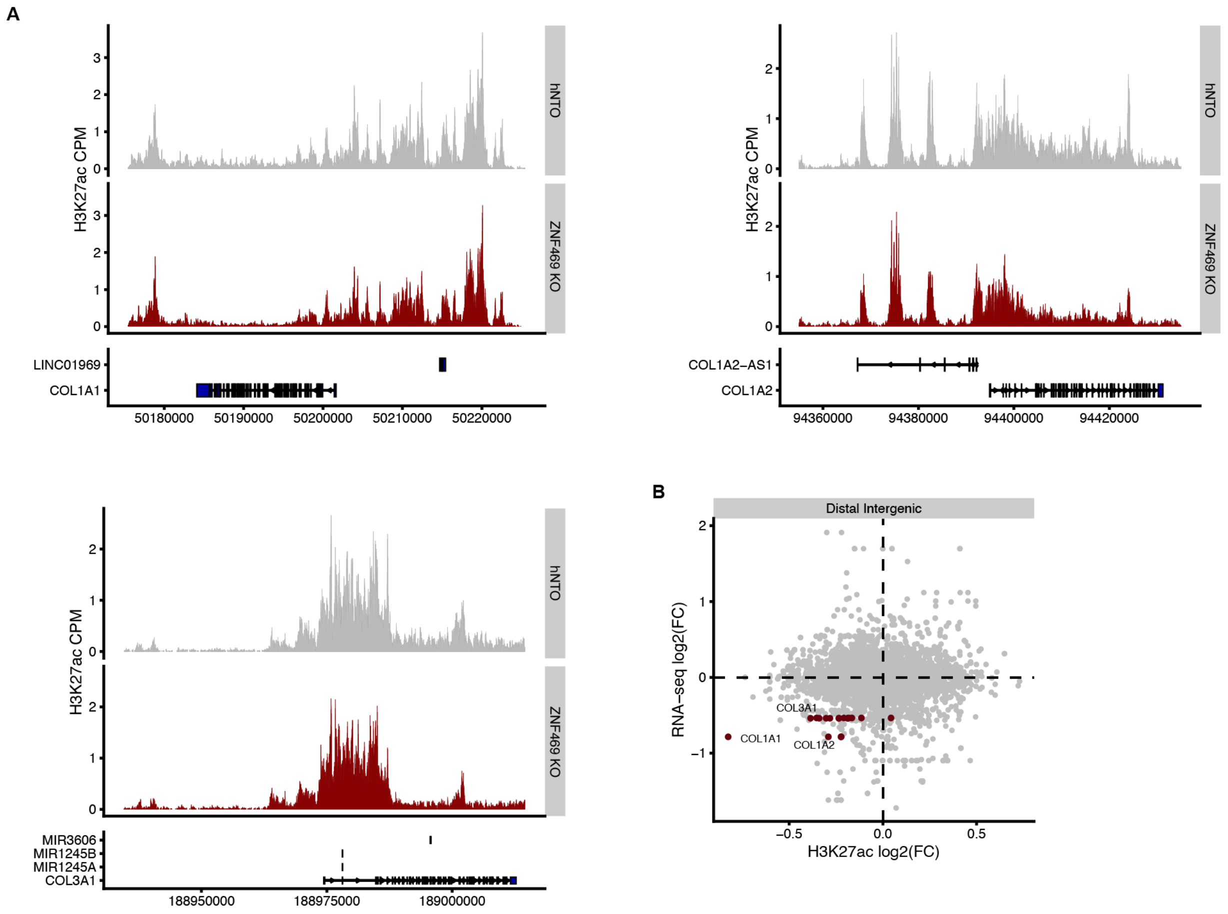


#### Supplementary Fig 5:

(**A**) Genome browser showing H3K27ac with or without *ZNF469* KO at *COL1A1*, *COL1A2* and *COL3A1* loci, (**B**) RNAseq and CUT&RUN integration upon *ZNF469* KO highlighting *COL1A1* and *COL1A2* as the top affected genes using distal intergenic regions.

Supp Figure 6: ZNF469 expression correlates with collagen production in HSCs in human NAFLD.

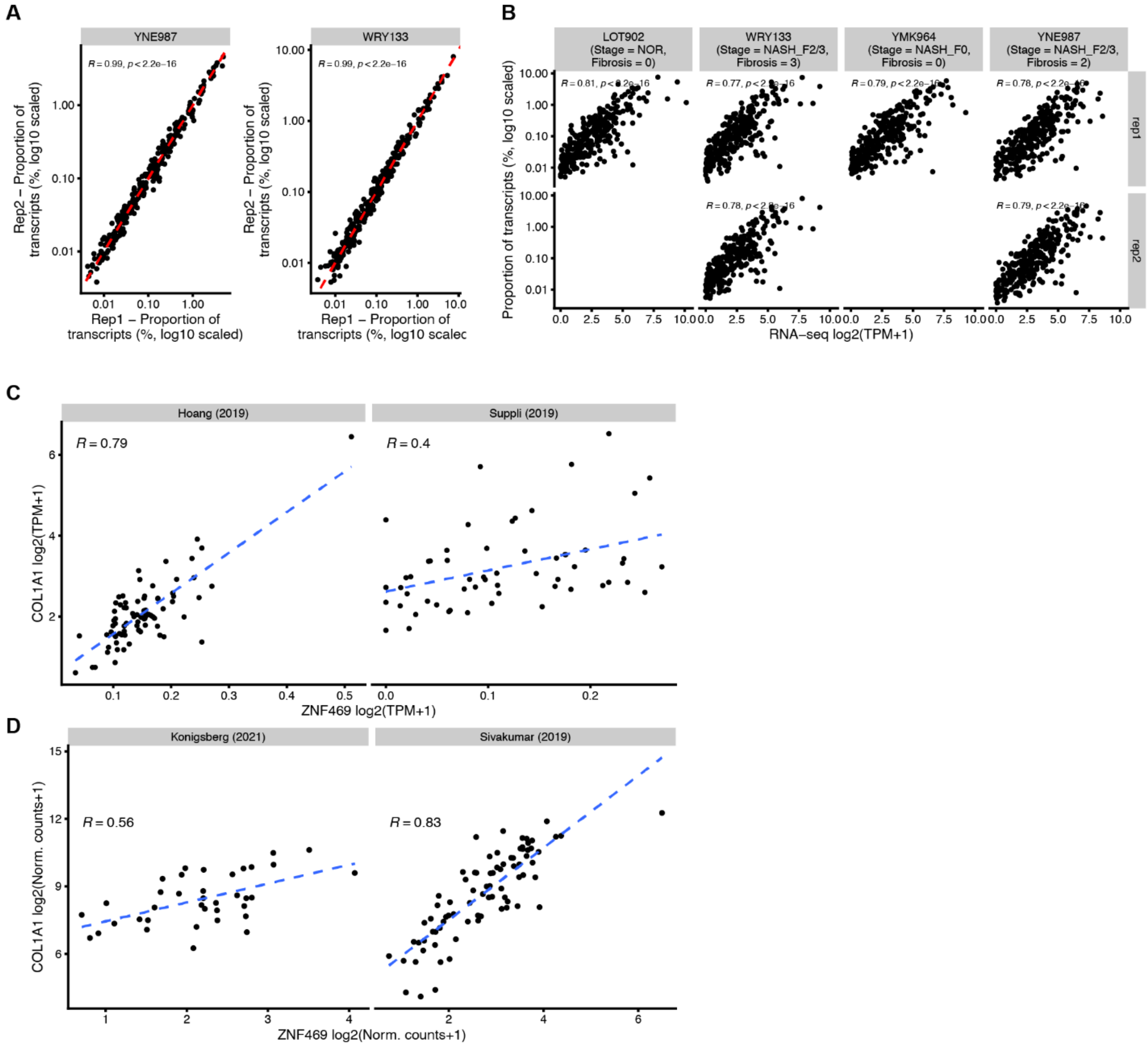


#### Supplementary Data Fig. 6:

(**A**) MERSCOPE correlation between replicates (**B**) MERSCOPE correlation compared to RNAseq (**C**) Scatter plot showing correlation (Pearson) of *ZNF469* and *COL1A1* expression in public NASH cohorts. (**D**) Scatter plot showing correlation (Pearson) of *ZNF469* and *COL1A1* expression in public interstitial pulmonary fibrosis cohorts.

**Supp Figure 7: ZNF469 expression correlates with collagen production in HSCs in mouse.**

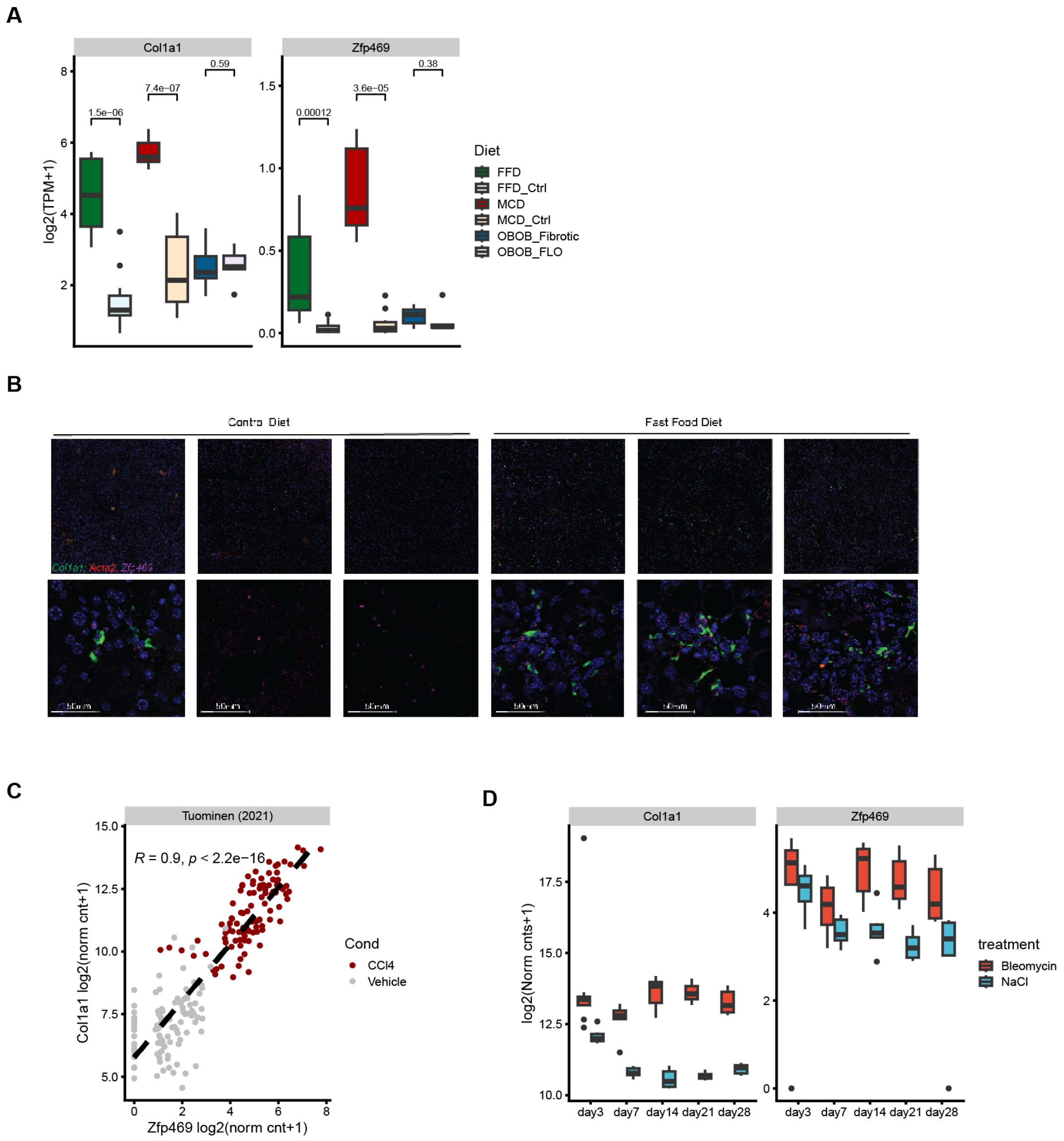


#### Supplementary Data Fig. 7:

(**A**) Box plots of relative differences in *Zfp469* mRNA RNA-seq datasets across three different mouse models of liver fibrosis (CDAHFD, GAN diet, ethionine-treated ob/ob). (**B**) RNAscope of *Zfp469* and *Col1a1* co-expression in control diet and GAN diet mouse livers. (**C**) *Zfp46*9 and *Col1a1* expression analyzed from publicly available CCL4 mouse model (liver), (**D**) *Zfp469* and *Col1a1* expression analyzed from publicly available bleomycin mouse model (lung).
